# Peptide Avidity for TCR on CD4 Effectors Determines the Extent of Memory Generation

**DOI:** 10.1101/2022.05.09.491158

**Authors:** Michael C. Jones, Catherine Castonguay, Padma P. Nanaware, Grant C. Weaver, Brian Stadinski, Olivia A. Kugler-Umana, Eric Huseby, Lawrence J. Stern, Priyadharshini Devarajan, Susan L. Swain

## Abstract

The impact of initial peptide antigen affinity for TCR in driving memory fate has been studied previously, however its contributions when effectors contract to memory is unclear. To become memory, effector CD4 T cells must recognize antigen at 5-8 days post-infection, at what we call the “effector checkpoint.” We examined whether peptide affinity for the TCR of effectors impacts the extent of memory and degree of protection against rechallenge. We made an influenza A virus (IAV) nucleoprotein (NP)-specific TCR transgenic strain, FluNP, and generated NP-peptide variants that bind FluNP TCR with a broad range of avidity. To vary avidity *in vivo*, we primed naïve donor FluNP in IAV-infected hosts, purified 6d FluNP effectors and co-transferred them with peptide-pulsed APC into uninfected second hosts. Higher affinity peptides yielded higher numbers of FluNP memory cells in the spleen and most dramatically in the lung and dLN, and drove better protection against lethal influenza infection. The major impact of avidity was on memory cell number, not cytokine production, and was already apparent within several days of transfer. We previously showed that autocrine IL-2 production during the effector checkpoint prevented default effector apoptosis and supported memory formation. Here, peptide avidity determined the level of IL-2 produced by effectors. IL-2Rα expression by APC drove more memory cell formation, suggesting that transpresentation of IL-2 by APC at this checkpoint enhanced CD4 memory generation. Secondary memory generation was also avidity-dependent. We propose this pathway selects CD4 effectors of highest affinity to progress to memory.

## Introduction

T cells recognize conserved viral epitopes and hence T cell memory provides broad, heterologous immunity crucial to protection against mutating viruses. Resting CD4 T cells respond to infection by generating effectors that protect against infection, via various synergizing mechanisms (1), including help to CD8 and B cells, production of IFNγ and perforin-mediated cytotoxicity. After viral clearance, a cohort of CD4 effectors becomes memory cells which can protect against future infections. However, the signals and mechanisms required for the transition of CD4 effectors to memory are only partially defined.

Several studies indicate that longer duration of Ag stimulation during priming results in an increased naïve CD4 T cell response, leading to increased proliferation and function of effector cells (2–4). Both the amount of peptide Ag loaded onto MHC-II receptors on an antigen presenting cell (APC), which determines the density of peptide-MHC (pMHC-II) complexes on the APC, and the affinity of the pMHC-II interaction with TCR, determine the extent of T cell response (5). While strong pMHC-II interactions with TCR favor Th1 over Th2 development (6–8), their role in T_FH_ versus Th1 differentiation is less clear, with conflicting reports (9–12). There are conflicting reports indicating either strong or weak initial pMHC-TCR interactions better support memory formation and recall responses (11, 13–20). Thus, whether CD4 effector cells transition to memory on the basis of their affinity for Ag is unclear.

Mice infected with influenza present a wide diversity of Ag epitopes to T cells, that reach very high levels soon after infection and that remain high until infection is cleared (21, 22). Infection produces very strong CD4 T cell memory, suggesting that persistent high levels of Ag, including some epitopes with high affinity, may explain the high levels of memory generated by infection. Here, we specifically analyze the impact of peptide avidity for TCR at the effector phase, when both viral levels and effectors have peaked, and when effectors that fail to recognize Ag undergo contraction (23, 24). We ask if high vs. low affinity Ag drives generation of more CD4 memory cells that provide superior protection. Previously, we showed that CD4 effectors generated *in situ* by IAV infection need to recognize Ag during the effector phase checkpoint, 5-8 days post infection (dpi), to become memory cells (23, 24). The effector cells must engage in a second cognate interaction and produce autocrine IL-2, which prevents their apoptosis, enabling their transition to memory (23–25). We ask here if the strength of p/MHC-TCR interaction plays a decisive role in driving CD4 effectors to memory.

To study the impact of peptide avidity at this crucial juncture, we developed a TCR Tg mouse (FluNP) specific for NP_311-325_, an immunodominant, highly conserved, IAV nucleoprotein (NP) epitope in B6 mice (22). We made a truncation and single amino acid substitutions to generate a library of NP_311_ peptides with a spectrum of functional avidities for the FluNP TCR. We generated *in vivo* effectors from naïve CD4 by IAV infection in a first (1^st^) host, isolated 6 dpi donor effectors and co-transferred them to a second (2^nd^) host (23) with peptides from a panel spanning high to low avidity for the FluNP TCR, as the only source of Ag.

We find that higher avidity Ag at day 6 (6d) of the CD4 effector response promotes a far larger CD4 memory population in the lung and dLN, and this leads to better protection. The peptide affinity and dose used to pulse APC determines the level of IL-2 produced by 6d effectors, and levels of IL-2 correlate with prevention of default apoptosis, effector cell survival and development of memory (24). CD25, the high affinity IL-2 receptor (IL-2Rα), is not expressed on 6d CD4 effectors, but is highly upregulated on APC in IAV-infected 4-8 dpi mice. Holding the peptide Ag constant, APC expressing CD25 generated higher numbers of CD4 T cell memory than CD25 deficient APC. We propose that IL-2Rα expression on APC acts in concert with IL-2Rβ/γ on the effector CD4 cells, to enhance local IL-2 signaling. We suggest at the effector phase, the level of IL-2 production, determined by peptide affinity, and efficacy of the response to autocrine IL-2, are the dominant pathways that regulate the size of the memory CD4 population and the extent of protection. We discuss the implications of this requirement for high peptide affinity at the effector stage for vaccine design with a focus on kinetics, Ag breadth and adjuvants that activate APC.

## Materials and Methods

### Mice

We use 8-12 wk old C57BL/6 (B6) mice as hosts in all experiments. Naive CD4 T cells are isolated from B6.FluNP strains, including B6.FluNP.Thy1.1^+/-^ and B6.FluNP.Nr4a1^EGFP^.Thy1.1^+/-^. The B6.Nr4a1 ^EGFP^ developed by Kris Hogquist and Steve Jameson (26) were from Jackson Laboratories. B6.FluNP TCR Tg mice were generated in collaboration with Eric Huseby’s Laboratory at UMMS. Briefly, B6 mice were infected with a sub-lethal dose of PR8 (0.3 LD_50_) and at 21 dpi, 2 x 10^7^ spleen and lung draining lymph node (dLN) cells were isolated and stimulated *in vitro* with irradiated spleen cells loaded with 100 ug/ml NP_311-325_ peptide. After 5 days, responding T cells were fused with BW5147 to generate T cell hybridomas (27). T cell hybridomas with reactivity to NP_311-325_ peptide, presented by lung APC from IAV-infected mice (A/PR8/34)-infected mice, were expanded. TCR Vβ-chains were identified by staining with a set of Vβ-specific Abs (BD Biosciences), and the TCRα-chains were identified by PCR analysis using a panel of TCR Vα primers that collectively amplify all TCR Vα gene families. We choose a hybridoma with Vα4.2 and Vβ2.1. A TCR Tg plasmid was made using cloned rearranged cDNAs for 22.B6 TCR Vα4.2 and Vβ2.1. Cloned products were fused with full length TCR Cα and Cβ sequences (28). All the TCR genes were sequenced, and error-free full-length cDNAs were subcloned into the human CD2 promoter transgene cassette for T cell-specific expression (29). B6.FluNP were established by injecting C57BL/6 oocytes with the TCR-Tg plasmid. BMDC were derived from B6 or B6.129S4-*Il2ra^tm1Dw^*/J (CD25KO) mice obtained from The Jackson Laboratory and bred at UMMS breading facility. Mice used in experiments were 8-12 wk of age. The Institutional Animal Care and Use Committee of UMMS approved all animal procedures.

### Virus Stocks and Infections

Mice were anesthetized with either isoflurane (Piramal Healthcare) or ketamine/xylazine (at a dose of 25/2.5 mg/kg by i.p. injection) before i.n. infection with 50 μl of influenza virus diluted in PBS corresponding to a 0.2 to 0.3 (sub-lethal) medial lethal dose (LD_50_) for response, 2LD_50_ for weight loss, and 4LD_50_ for survival. Influenza virus A/Puerto Rico/8/34 (PR8, H1N1), originally from St. Jude Children’s Hospital, was from our stocks grown and maintained at the Trudeau Institute. The virus was also characterized by its ability to infect eggs, and we found 2LD_50_ corresponds about 10,000 EID_50_. Our standard dose sub-lethal 0.3LD_50_, corresponding to 25 PFU.

### NP Peptide Generation

We modified the NP_311-325_ peptide to produce peptides of shorter lengths by deletions of amino acids (aa) on both ends to determine the best length. We used single alanine substitutions to determine the peptide-I-A^b^ binding frame (P1=**Y**), then selective aa side-chain modifications, at known peptide-TCR contact positions to generate peptides likely to have lower affinities. Peptide-I-A^b^ IC_50_ was determined with surface plasmon resonance (SPR) using a BIAcore 3000. SPR analysis was performed using a BIAcore 3000 instrument (Cytiva). Briefly, biotinylated, peptide exchanged MHCs were immobilized on a streptavidin chip. For the TCR, the FluNP TCR sequence was cloned into the pCDH lentiviral expression vector with a P2A site separating the alpha and beta chains. This construct was used to generate stable lines in 293S GnTI cells. TCR was then purified from supernatant with a nickel-NTA column and subsequent size exclusion. Recombinant FluNP TCR was then passed over at increasing concentrations. A5 was used as a negative control and the signal from this flow cell was subtracted from those of experimental flow cells. The resulting data points were plotted and fitted to hyperbolas to derive K_D_s.

### BMDC Generation and Peptide/APC Preparation

APC were generated as in (23, 30) BM was harvested from B6 or CD25KO mice, washed with RPMI 1640, 1% FBS and cells plated at10^7^ cells/mL in RPMI with 10% FBS and 10 ng/mL GM-CSF. After 7d, CD11c^+^ BMDC were isolated via MACS and activated with polyinosinic-polycytidylic acid (PolyI:C) at 10 μg/mL overnight in culture or use as APC. APC were pulsed with a standard concentration of 100 μM or dilutions thereof, of each of a chosen panel of NP peptides. For *in vivo* experiments pulsing was at 37°C for 1hr with shaking, APC were washed 3x, resuspended in PBS, and 1 x 10^6^ cells injected i.v. per mouse.

### Sequential Transfer Model

We closely followed the model we developed previously (23). Spleens and peripheral lymph nodes (LN) were collected from B6.FluNP.Thy1.1^+/-^ mice. Naïve CD4 cells were isolated via negative selection with CD4 MACS (Miltenyi Biotec and washed 3x, resuspended in PBS. 0.5-1 x 10^6^ naïve CD4 cells were transferred via i.v. injection into B6 1^st^ hosts. 1^st^ hosts were infected with a sub-lethal dose of PR8 the same day. Donor CD4 effectors were re-isolated from the 1^st^ hosts at 6 dpi. Single cell suspensions were prepared from pooled spleen and dLN and donor FluNP cells were isolated via Thy1.1 positive selection by MACS (Miltenyi Biotec). Cells were resuspended in PBS and 1.5 x 10^6^ donor FluNP cells per mouse injected into 2^nd^ B6 hosts i.v., along with Ag/APC (peptide-pulsed). To maintain effector phenotype all steps were conducted at room temperature, except for one 15 min incubation at 4°C. This minimal protocol, without sorting, ensures that effector cells are only out of mice for a maximum of 2.5 hours

### *In Vitro* Culture of Naïve and 6 dpi effector FluNP CD4 T cells

To assess functional avidity of peptides, we compared their ability to induce responses of FluNP naïve CD4 T cells, isolated as described above (sequential transfer model). Following isolation of naïve or 6d FluNP effectors and generation of Ag/APC, cells in complete RPMI 1640 media were plated at a CD4 T:APC ratio of 5:1. Following 2d in culture, plates were centrifuged, supernatant was removed for cytokine protein analysis via ELISA and cell pellets were stained for FACS analysis.

### ELISA

Supernatants were collected from *in vitro* culture. Plates (Nunc) were coated overnight with capture antibody (ELISAmax Biolegend). The following day plates were blocked following the manufacturer’s protocol and supernatants were added neat, or diluted 1:10, 1:100, 1:1000 and left at 4°C overnight. The following day plates were washed, and detection protocol was followed per manufacturer’s protocol.

### Flow Cytometry and Cytokine Staining

Cells were harvested, passed through a 70 μm cell strainer and stained in FACS buffer (0.5% Bovine Serum Albumin, 0.01% sodium azide (Sigma Aldrich) in PBS. Cells were blocked with anti-FcR (2.4G2) and stained with amine reactive viability dyes (Invitrogen) to exclude dead cells. Surface proteins were stained with fluorochrome conjugated antibodies at 4°C. Antibodies used included anti: CD4 (GK1.5 and RM), CD44 (IM7), CD90.1 (OX-7 and HIS51), CD11c, CD25, CD62L, CD69, CD80, CD86, CD122, CD132, CD185 (CXCR5, SPRCL5), NKG2A/C/E, MHCII (I-A^b^). For CD127 staining, anti-CD127-Biotin was included in the surface stain mixture, cells were washed 3x, and a secondary fluorochrome conjugated SA was used in the second step. For cytokine staining, total splenocytes were stimulated with 10 μM of NP_311-325_ for 6 hr at 37°C. Brefeldin A (10 μg/mL) was added after 1 hr of stimulation. Following surface staining, cells were fixed in 2% paraformaldehyde for 20 min and permeabilized with 0.1% saponin buffer (1% FBS, 0.1% NaN_3_ and 0.1% saponin in PBS (Sigma Aldrich) for 15 min. Subsequent staining for cytokines using the following antibodies: anti-IFNγ (XMG1.2), anti-TNFα (MP6-XT22), anti-IL-2, anti-IL-17. For transcription factor staining cells were first surface stained then fixed and permeabilized using FoxP3 fix/perm kit (eBioscience) overnight per manufacturer’s protocol and stained with the following antibodies: anti-BCL-6 (K112-91), anti-FoxP3, anti-T-bet at 4°C for 1 hour. Antibodies obtained from BD Bioscience, Biolegend and eBiosceince. Stained cells were acquired on a BD LSRII flow cytometer and analyzed using FlowJo analysis software.

### Statistical Analysis

Groups of at 3-5 mice were used in all experiments, and exact conditions repeated to obtain sufficient statistical power. All experiments shown were repeated 2-3 or more times. For statistical analysis an unpaired, two-tailed independent t test was used. All analysis was performed using GraphPad’s Prism.

## Results

### Characterization of FluNP TCR transgenic CD4 T cell Response

We generated a B6 TCR transgenic (Tg) mouse specific for an immunodominant NP core protein epitope, NP_311-325_, from the internal nucleoprotein (NP) of influenza PR8/34 (H1N1) presented by I-A^b^ (MHC-II). This epitope is conserved among all dominant outbreak strains of IAV in people (22). The mouse was created by selecting T cell hybridomas specific for NP_311-325_ from IAV-infected mice (27–29). We call the mouse FluNP.

We generated B6.FluNP.Thy1.1/Thy1.2 mice so we could readily detect donor FluNP cells after transfer to new hosts using Thy1.1 expression. To evaluate whether IAV induces a comparable response of the donor FluNP cells and polyclonal host CD4 T cells, we transferred naïve FluNP CD4 T cells into hosts and infected with a sub-lethal dose of PR8/34, an influenza A virus (IAV). We compared the kinetics of donor FluNP TCR Tg and endogenous host (CD4^+^CD44^hi^) T cell responses 4, 6, 8, 12, 21 and 63 days post infection (dpi) in the lung (Fig. 1A), draining mediastinal lymph node (dLN) and spleen (Supp. Fig. 1A). IAV infection induced a similar pattern of expansion and contraction of donor FluNP and host CD4 T cells in the lung (Fig. 1A), spleen and dLN (Supp. Fig. 1A). To evaluate the subsets of CD4 effectors generated, we compared donor and host CD4 effectors at 8 dpi by examining cytokine production and phenotype markers: Th1 (T-bet, IFNγ, TNFα, IL-2), Triple cytokine producers (IFNγ, TNFα, IL-2), Th17 (IL-17), and T_REG_ (FoxP3). We used NKG2A/C/E expression to detect cytotoxic CD4 (ThCTL), found only in the lung (31), and CXCR5 and Bcl-6 co-expression markers for T_FH_ in the spleen (32) (Fig. 1B, Supp. Fig. 1B-I). Both donor and host responses in the lung were dominated by Th1 phenotype cells (33–35),with high expression of T-bet, IFNγ and TNFα (Fig. 1B, left, Supp. Fig. 1B, E), and little expression of IL-17 or FoxP3 (Fig. 1B, left, Supp. Fig. 1D). ThCTL were found in both at similar proportions (Fig 1B, left, Supp. Fig. 1C). In the spleen, donor and host subset patterns were also similar (Fig 1B, right, Supp. Fig. 1F-I). Overall, the effector responses of donor monoclonal FluNP and host polyclonal CD4 to IAV were comparable.

**Figure 1:**
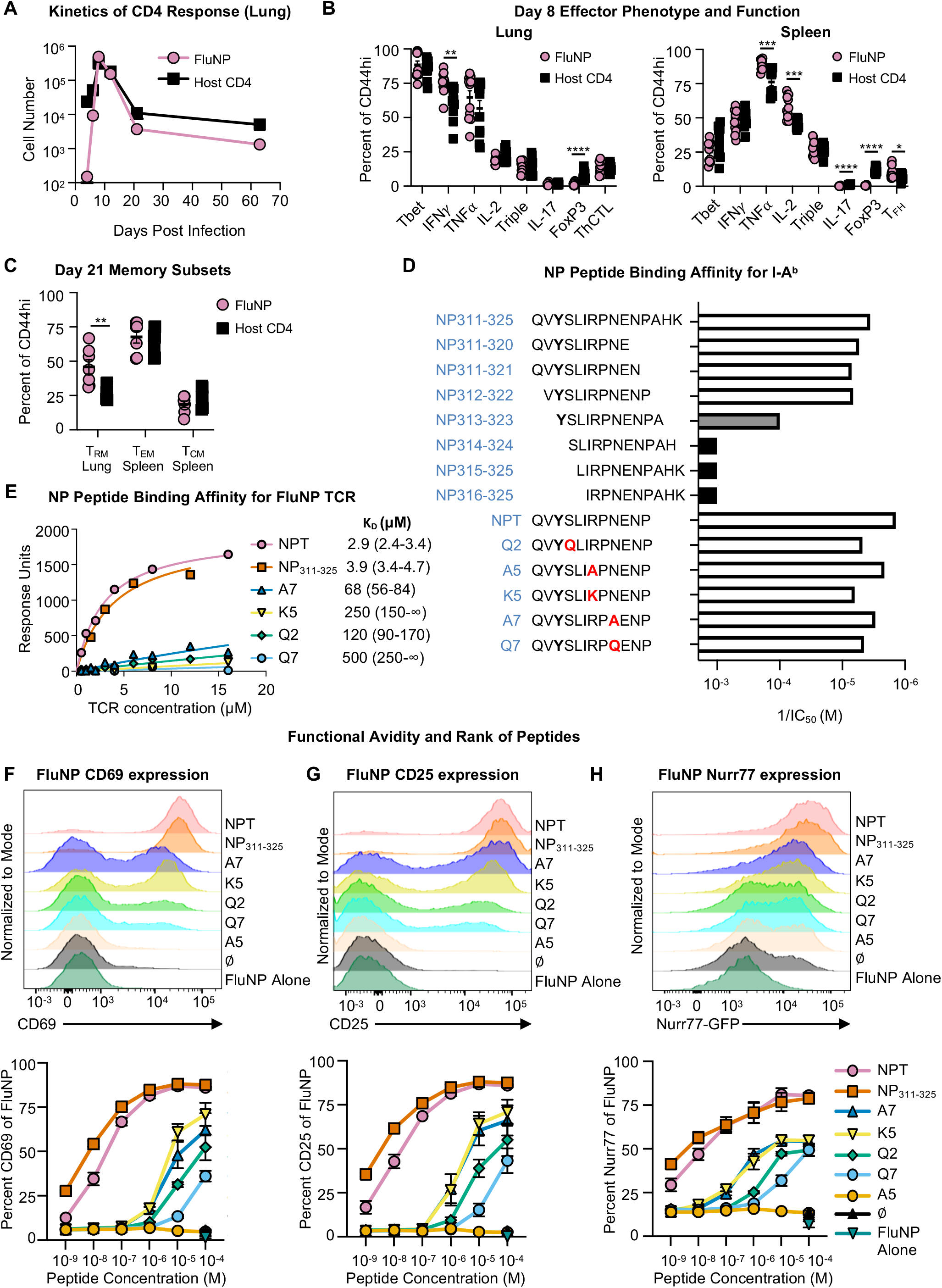
Model to evaluate peptide avidity. **(A-C)** Naïve FluNP.Thy1.1^+/-^ cells were transferred to B6 hosts, then infected with PR8. **(A)** Lungs were collected at 4, 6, 8, 12, 21 and 63 dpi and numbers of donor FluNP and responding host CD4^+^, CD44^hi^ cells were determined by FACS. **(B)** Day 8 Effector phenotype and function. Expression of markers associated with subsets of CD4 effectors were analyzed at 8 dpi in lung and spleen: Th1 (T-Bet^+^, IL-2^+^, IFNγ^+^, TNFα^+^), Triple positive (IL-2^+^, IFNγ^+^, TNFα^+^), Th17 (IL-17^+^), Treg (FoxP3^+^), lung ThCTL (NKG2A/C/E) and spleen T_FH_ (CXCR5^+^, BCL-6^+^). **(C)** Memory CD4 subsets were analyzed at 21 dpi: lung T_RM_ (CD69^+^), spleen T_EM_ (CD127^+^ CD44^+^ CD62L^-^) and spleen T_CM_ (CD127^+^CD44^+^ CD62L^+^). **(D-H)** Characterization of NP peptide panel. **(D)** Peptide name, aa sequence (P1=**Y**, mutations in red) and the I-A^b^ binding affinities of the NP_311-325_ length variants and mutants were experimentally determined to identify the I-A^b^ binding frame. The reciprocal of the IC_50_ is shown. **(E)** Maximal response for a given TCR concentration was plotted for each peptide-MHC complex and nonlinear fits were generated using the equation Y=Bmax*X/(KD + X). The fit was constrained to share a consistent Bmax and yield a K_D_ value of <500. The resulting K_D_ values are shown in μM with 95% confidence intervals. **(F-H)** Naïve FluNP CD4 T cells were co-cultured with BMDC pulsed with each of the NP peptides for 2d *in vitro.* Induction of markers functionally associated with TCR signal strength was measured. **(F)** CD69; **(G)** CD25; and **(H)** Nur77. Top histograms display level of marker expression following stimulation with Ag/APC pulsed at 10^-4^ M. Bottom displays dose response curve to a broad range of peptide concentrations used to pulse APC. The rank of peptide functional avidity is shown on right. Statistical evaluations: (A) Days 4, 6, 8, 21, 63 Pooled data, n = 9-10, two experiments. Day 12 one experiment n = 5. Mean +/- SEM. (B) Pooled data, n = 11, two experiments, mean +/-SEM. (C) Pooled data, n = 7, two experiments, mean +/- SEM. (D) Pooled data, n=3-4, two experiments. (E-G) Representative data, n = 6, two experiments. (F-H) Pooled data, n = 6, two experiments, mean +/- SEM (% of FluNP). Statistical significance determined by two-tailed independent t test (*p<0.05, **p<0.01, ***p<0.001, ****p<0.0001).

We examined memory populations at 21 dpi, focusing on phenotypically distinct memory subsets: central memory T_CM_ (CD127^+^ CD44^+^ CD62L^+^), effector memory T_EM_ (CD127^+^ CD44^+^CD62L^-^) and resident memory T_RM_ (CD44^+^ CD69^+^). The overall patterns in donor and host memory were again similar, (Fig. 1C, Supp. Fig. 1J-M) and both donor and host memory cells expressed high levels of the canonical CD4 memory marker CD127 (~75%), in all tissues (Supp. Fig 1L). This supports the suitability of this model to investigate the impact of peptide avidity on memory generation.

### Generation of an NP Peptide Panel Spanning a Range of Avidities and Affinities for FluNP TCR

To study the role of peptide avidity, we generated a series of NP peptides with single amino acid (aa) substitutions that sample a broad range of abilities to stimulate the naïve FluNP response when pulsed on B6-derived activated APC. We sought to identify substitutions that altered TCR interaction while maintaining tight peptide-MHC interaction. First, we identified the I-A^b^ binding frame by scanning partially overlapping 11-residue peptides that cover the full NP_311-325_ peptide for I-A^b^ binding, using a fluorescent peptide competition binding assay and purified recombinant I-A^b^ carrying a cleavable linker peptide (36). I-A^b^ binding was substantially reduced for NP_313-323_ and lost completely for NP_314-324_, (Fig. 1D) suggesting that Y_313_ occupied the key P1 position in the I-A^b^ binding site (37, 38). We confirmed this using alanine-scanning mutagenesis, revealing Y_313_ as the only position where I-A^b^ binding was substantially affected (Fig. 1D). To identify peptides that modulate TCR interaction, we introduced other substitutions in addition to alanine at the predicted TCR contact positions P2 (Ser→Gln), P5 (Arg→Lys), and P7 (Glu→Gln). Some of these substitutions caused moderate reductions in peptide-MHC binding, up to 3.4 for Q2 (Fig. 1D). To evaluate the effect of these substitutions on FluNP TCR interaction independent of peptide-MHC effects, we measured pMHC-TCR binding using a surface plasmon resonance (BIAcore) assay with streptavidin-immobilized biotinylated I-A^b^-peptide complexes and recombinant soluble FluNP TCR (Fig. 1E). FluNP TCR bound to I-A^b^ carrying the parent NP_311-325_ peptide or truncated NP_311-322_ (NPT) with high affinity (K_D_ ~ 3uM) (Fig. 1E). The other substitutions at predicted TCR contact position caused reductions in pMHC-TCR affinity ranging from ~20-fold (for A7) to >150-fold (for Q7). The A5 substitution abrogated detectable binding.

To rank the substituted NP peptides by their ability to stimulate a FluNP response (functional avidity), we loaded bone marrow derived dendritic cells (BMDC) with peptides over a broad dose range and evaluated how well they stimulated naïve FluNP cells *in vitro*. After 2d, we assessed induction of CD69 (Fig. 1F), CD25 (IL-2Rα) (Fig. 1G) and Nurr77 (Fig. 1H). By all three assays, FluNP CD4 T cells responded in a dose and affinity dependent manner. Thus, we could confidently rank the relative avidity of the peptide-MHC-II complex on APC for the FluNP TCR on the CD4 T cells. We classify the NPT & NP_311-325_ as high avidity, A7 & K5 as medium (mid) avidity, Q2 & Q7 as low avidity, and A5 and unpulsed BMDC as negative controls (neg). In each assay, the high peptides (NP_311_ and NPT) induce peak responses at doses 100-fold lower than the middle peptides (A7, K5), and the low peptides (Q2, Q7) require 10 times the dose as the two middle peptides (Fig. 1F-H).

From here on, we use a high dose of 10^-4^ M to pulse APC, to minimize the contribution of peptide density and maximize the contribution of pMHC-TCR affinity. At this concentration, all 6 peptides (NPT, NP_311-325_, A7, K5, Q2, Q7) stimulate a measurable FluNP naïve CD4 T cell response.

### Peptide Avidity at the Effector Phase determines the Number of Memory Cells

We evaluated the impact of peptide avidity on *in vivo* memory generation using a sequential adoptive transfer model, developed previously (23). *In vivo* IAV generated 6d FluNP effectors were co-transferred to uninfected 2^nd^ hosts with groups of peptide-pulsed APC as the only source of Ag (23) (Fig 2C). The APC are short-lived and present Ag for only 48-72h (23), defining the discrete checkpoint of Ag recognition. We evaluated how the magnitude of FluNP effector response to high avidity peptide/APC compares to a polyclonal response to IAV (Fig. 2A). At 21 dpi, NPT/APC stimulated FluNP effectors produced as many memory cells in spleen and lung and only slightly fewer in dLN as did PR8 infection (Fig. 2B). Thus, a high avidity peptide/APC generates an equivalent number of memory cells from effectors as does IAV infection as seen in an equivalent model using OT-II Tg CD4 T cells (23).

**Figure 2:**
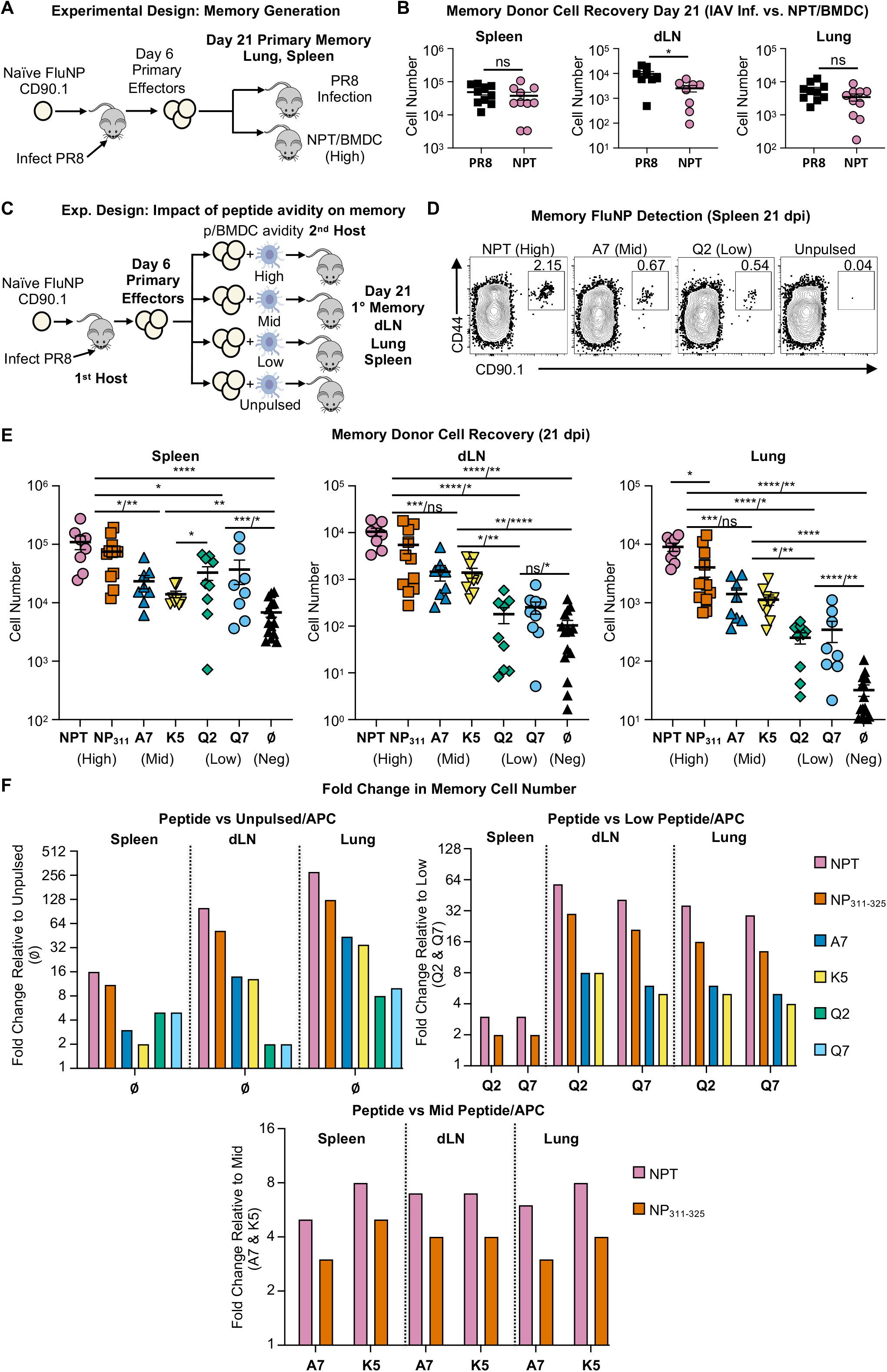
Peptide avidity during the primary effector phase dictates the size of the memory population. **(A)** Experimental design: comparison of PR8 infection to high peptide/APC memory generation. Naïve FluNP.Thy1.1^+/-^ cells were transferred to B6 hosts, then infected with PR8. At 6 dpi, FluNP effector cells were isolated from the 1^st^ hosts and co-transferred with either peptide Ag/APC into uninfected 2^nd^ hosts or without APC into day 6 PR8 infection matched hosts (infected 6d previously). Fifteen days later (21 dpi) 2^nd^ hosts were sacrificed, and donor FluNP cells were analyzed by FACS. **(B)** FluNP cell numbers were enumerated by FACS in the dLN, lung and spleen at 21 dpi. **(C)** Experimental design: Impact of peptide avidity on memory generation. 1.5×10^6^ 6d FluNP effectors were co-transferred with Ag/APC into uninfected 2^nd^ hosts. The panel of different peptides was used to pulse groups of activated BMDC yielding peptide/APC with different avidity for the TCR of the FluNP T cells. **(D)** Representative FACS plots showing donor FluNP memory cells by CD44 and CD90.1 expression in the spleen at 21 dpi. **(E)** Number of FluNP memory cells detected at 21 dpi in the dLN, lung and spleen of second hosts. Statistical significance determined by two-tailed independent t test (*p<0.05, **p<0.01, ***p<0.001, ****p<0.0001). <u>Spleen comparisons</u>: highs vs. unpulsed, highs vs. lows, NPT vs. A7 / highs vs. K5; mids vs. unpulsed, K5 vs. Q2; Q2 vs. unpulsed / Q7 vs. unpulsed. <u>dLN comparisons</u>: NPT vs. unpulsed / NP_311-325_ vs. unpulsed, NPT vs. lows / NP_311-325_ vs. lows, NPT vs. mids / NP_311-325_ vs. mids; A7 vs. unpulsed / K5 vs. unpulsed, A7 vs. lows / K5 vs. lows; Q2 vs. unpulsed / Q7 vs. unpulsed. <u>Lung comparisons</u>: NPT vs. NP_311-325_; NPT vs. unpulsed / NP_311-325_ vs. unpulsed, NPT vs. lows / NP_311-325_ vs. lows, NPT vs. mids / NP_311-325_ vs. mids; mids vs. unpulsed, mids vs. Q2 / mids vs. Q7; Q2 vs. unpulsed / Q7 vs. unpulsed. **(F)** FluNP cell number fold change relative to unpulsed (left), low peptide-pulsed (right) and mid peptide (below) in dLN, lung and spleen 21 dpi, x-axis lists fold change denominator. (B) Pooled data, n = 10, three experiments, mean +/- SEM. (D) Representative data, n = 8-15, four experiments. (E-F) Pooled data, n = 8-15, four experiments, mean +/- SEM.

We investigated if modulating avidity over a broad range would lead to a corresponding influence on donor memory numbers in the 2^nd^ hosts in the same model (Fig. 2C). The impact of peptide avidity was striking, with memory cell numbers clearly dependent on the rank of the peptides. The greatest differences in memory cell number were seen in the lung, followed by the dLN, and lastly the spleen (Fig. 2D-E, Supp. Fig. 2A). The significance of these changes between groups of peptides is illustrated by the fold change in memory cell number between each peptide compared to unpulsed (Fig. 2F, left), to low peptides (2F, right) and to middle peptides (Fig. 2F bottom). In the lung, there were 280-fold more donor FluNP cells in the NPT pulsed group compared to the unpulsed group, 29 and 36-fold more compared to the low groups and 6 and 8-fold more donor memory cells compared to the mid groups (Fig. 2F). A nearly identical pattern was seen in the dLN. In spleen, there were 16-fold more FluNP cells in the NPT group compared to the unpulsed group and 5-8-fold to the mid groups, both highly significant (Fig. 2E-F). These data indicate that the strength of pMHC-TCR interaction at the effector checkpoint determines the size of the memory population.

The donor memory cells uniformly express high CD127, the IL-7Rα which supports CD4 memory generation in the secondary lymphoid tissues (39), in all stimulated groups in the spleen and dLN (Fig 2 C-E), while in the lung many donor cells do not express CD127. We assessed cytokine production *ex vivo* by the memory cells stimulated with different peptides on APC by intracellular cytokine staining (ICCS) (Supp. Fig. 2F-H). Overall, all memory cells produced equivalent levels of IL-2, TNFα, and IFNγ. The high affinity peptide resulted in only a slightly higher fraction of IFNγ-capable memory. Overall, these results suggest the fewer memory cells that develop at lower avidity are likely to be functional.

### Peptide avidity at the effector checkpoint acts by promoting effector cell survival

To probe the mechanisms by which peptide avidity has such a dramatic impact on memory cell recovery, we asked if affinity impacts FluNP effector cell proliferation, survival, or both (Fig. 3A). To evaluate proliferation, we stained the 6d donor effectors with cell trace violet (CTV) before transfer and examined their division *in vivo* 3d after co-transfer with Ag/APC. As indicated by dilution of CTV, the spleen FluNP cells in each peptide group divided multiple times, while the unpulsed group barely proliferated (Fig. 3B). Thus, proliferation of transferred cells required Ag recognition, but peptide avidity did not impact the pace or extent of division in in any organ (Supp. Fig. 3A-C). Thus, the dramatic differences in memory cell numbers (Fig. 2), are not due to a greater rate or number of cell divisions.

**Figure 3:**
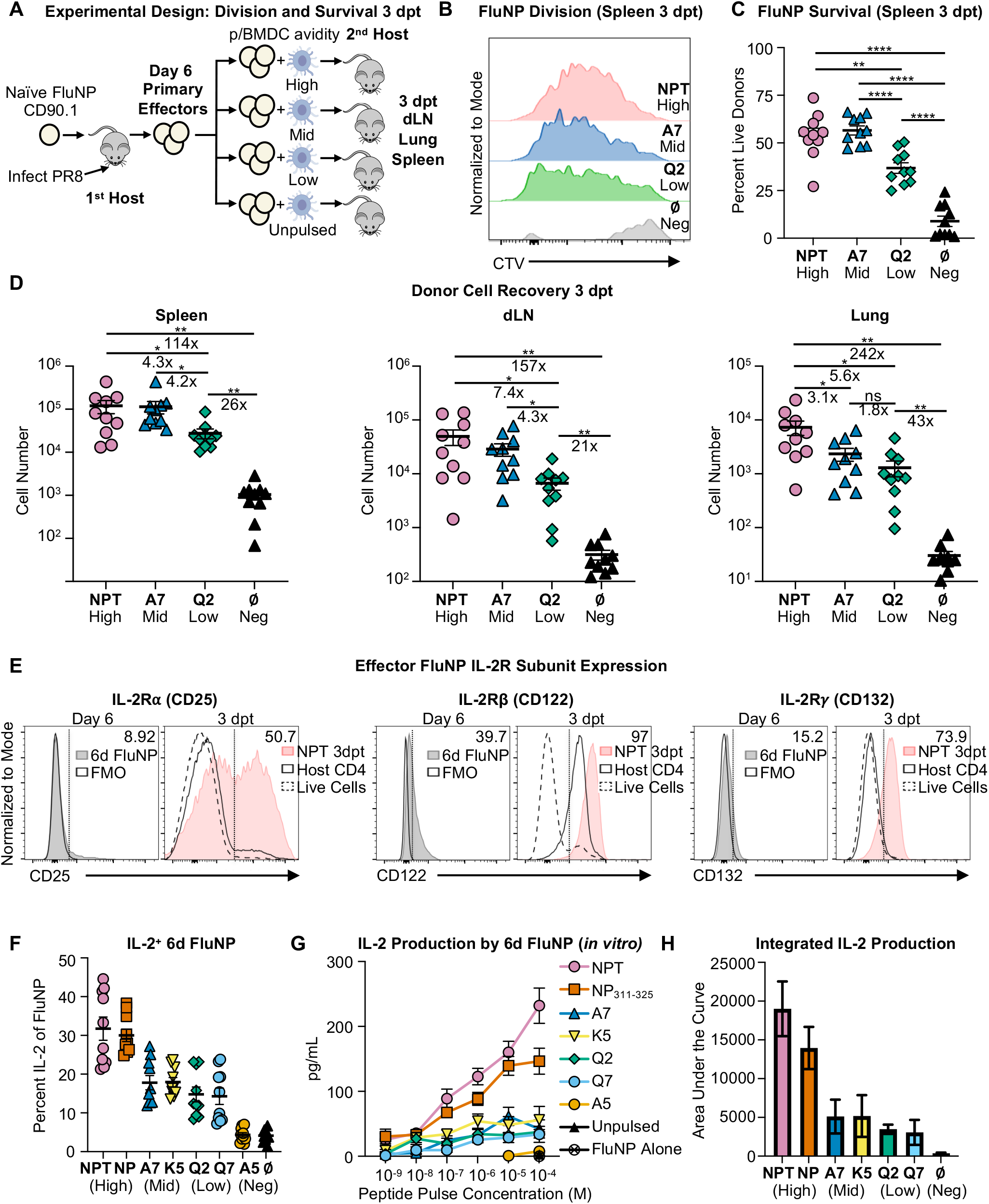
Peptide avidity determines survival but not proliferation of donor cells 3 days post transfer. **(A)** Experimental Design. Naïve FluNP CD4 T cells were transferred to B6 hosts, infected with PR8. At 6 dpi, FluNP effector cells were isolated from the 1^st^ host, labeled with cell trace violet (CTV) and transferred to 2^nd^ hosts given 10^6^ APC, pulsed or not, with high, mid or low peptides. At 3 days post transfer (dpt), 2^nd^ hosts were sacrificed and FluNP (CD4^+^ CD90.1^+^) cells were analyzed. **(B)** Representative FACS plots of FluNP CTV dilution in spleen 3 dpt. Gated on live singlets, CD4^+^, CD90.1^+^ cells. **(C)** Percent survival of FluNP effectors measured by live/dead and caspase 3/7 FACS (double-negative) in spleen 3 dpt. **(D)** Number of donor FluNP cells recovered 3 dpt in the dLN, lung and spleen. **(E)** IL-2 receptor subunit expression: IL-2Rα (CD25), IL-2Rβ (CD122) and IL-2Rγ (CD132) expression was determined by FACS analysis of the 6 dpi effectors (left) and 3 dpt donor cells (right). **(F)** FluNP 6 dpi effectors were restimulated for 6 hours with indicated NP peptides at 10 μM and IL-2 expression determined by FACS. **(G)** FluNP effectors were co-cultured with NP peptide-pulsed APC for 48 hr and IL-2 production in the supernatant was determined by ELISA. **(H)** The area under the curve was calculated to quantitate impact on IL-2 production. (B) Representative data, n = 10, two experiments. (C-E) Pooled data, n = 10, two experiments, mean +/- SEM. (F) Pooled data, n = 10, two experiments, mean +/- SEM. (G-H) Pooled data, n = 3-6, two experiments, mean +/- SEM.

We determined survival of FluNP effectors by measuring expression of caspase 3/7 and viability stain (Fig. 3C, Supp. Fig 3D-F). In the spleen, we found a much greater fraction of live, caspase 3/7 negative cells in the NPT (high) and A7 (mid) groups compared to the Q2 (low) group (Fig. 3C, Supp. Fig. 3D), indicating there is greater apoptosis in the low avidity group compared to both mid and high groups. Most donor cells recovered in the unpulsed APC group were pro-apoptotic by 3d post-transfer (dpt) (Fig. 3C), consistent with a strict requirement for Ag recognition to prevent apoptosis (23). We did not see a clear difference in cell survival with different affinity peptides in dLN or lung, although all peptides led to higher survival than no peptide (Supp. Fig 3E-F). Perhaps the response kinetics are different in each organ, or the spleen is a major source of memory generation after which developing memory migrate to the lung and dLN. Therefore, we also evaluated donor cell recovery as a measure of net survival. We found a clear-cut pattern with recovery corresponding to higher peptide avidity in all sites (Fig. 3D) that was of similar magnitude at 3 dpt (late effector) as it is at 15 dpt (memory) (Fig. 2E). We note that the impact of peptide avidity in regulating memory cell recovery is realized within a few days after effector cell Ag encounter, and it is primarily due to the fraction of effectors that survive rather than their extent of division.

### Peptide avidity at the effector checkpoint acts by inducing more effector IL-2 production

We established that effector phase autocrine IL-2 signaling is required for CD4 T cell memory and that autocrine IL-2 acts by downregulating pro-apoptotic Bim which promotes the survival of CD4 effectors (23, 24). We thought this implied that Ag recognition induced IL-2Rα expression as well, since anti-IL-2R blocked the impact of IL-2 (24). We analyzed the expression of IL-2R subunits CD132 (IL-2Rγ), CD122 (IL-2Rβ) and CD25 (IL-2Rα) on donor FluNP before transfer and 3d after *in vivo* stimulation with the various peptide/APC. Before restimulation, a few FluNP effectors express CD132 (15%), a fair fraction expresses low levels of CD122 (40%) but very few express CD25 (<10%) (Fig. 3E). Following stimulation with high affinity peptide/APC *in vivo*, three-quarters of FluNP cells in lung 3 dpt strongly express CD132 and almost all express CD122 (>90%) (Fig 3E), but this is true regardless of peptide avidity. Moreover, we found there are few, if any, FluNP effectors that express any CD25 in spleen and lung (Supp. Fig. 3G), and only half in the dLN (Fig 3E, Supp. Fig. 3G). In the *in vivo* setting, we found no difference in IL-2R subunit expression on 3 dpt donor cells following high, mid and low TCR signaling in any tissue, and even the unpulsed group 3 dpt effectors expressed a similar pattern (Supp. Fig. 3G), suggesting avidity does not act via IL-2R expression, leaving unresolved how autocrine IL-2 effectively signals the effectors.

We examined the impact of peptide avidity on the level of IL-2 produced by 6d effectors. We restimulated 6d FluNP effectors *ex vivo* with our panel of NP peptide-pulsed APC, and quantified IL-2-producing cells by ICCS (Fig. 3F) and levels of secreted IL-2 by ELISA (Fig. 3G-H). The fraction of IL-2-positive 6d FluNP effectors corresponded closely with relative peptide avidity, mirroring the impact on memory. Thus, induction of IL-2 secretion by FluNP effector cells is highly dependent on avidity of Ag recognition. Since the CD4 T cell response is also dependent on density of peptide Ag/MHC we varied the peptide concentration used to pulse APC (Fig. 3G, Supp. Fig. 3J). We found dose also strongly influenced IL-2 production by the 6d effectors. With high peptides, levels of IL-2 increased with concentration and were detectable even at very low dose (10^-7^ M) while mid and low peptides required much higher concentrations. We integrated the area under the curve, as a reflection of IL-2 accumulation (Fig 3H). This confirmed IL-2 production mirrored the rank of peptides determined by measures of affinity and functional avidity (Fig. 1). As peptide avidity determines the level of autocrine IL-2 produced, and since IL-2 availability determines their survival (24), this is likely the key pathway that translates peptide avidity to number of memory cells generated. IFNγ and TNFα production was also dependent on peptide avidity and on dose (Supp. Fig 3H-I), indicating the elicitation of those key effector function also is avidity driven

### CD25 expression on APC during the effector phase promotes effector transition to memory

When effector cells recognize peptide Ag presented by APC, they make IL-2 within several hours (40), which traditionally was predicted to bind to the tripartite IL-2R on the same cell, since the IL-2 needs to be autocrine (24). However, since 6d FluNP effectors do not express CD25 (Fig. 3E), we further investigated this mechanism. IL-15 shares two-thirds of its receptor (IL-2Rβ/γ) with IL-2 and is known to be transpresented by IL-15Rα on APC to IL-2Rβ/γ on T cells (41–43). Several previous studies found that IL-2 can also be transpresented (44–46). If IL-2 transpresentation plays a role here, IAV infection should generate CD25-expressing APC. We analyzed infection-induced APC for CD25 expression. In uninfected mice, we found no CD25 expression on MHC-II^+^ cells but at 6 dpi a cohort of dLN CD11c^+^, MHC-II^+^ cells clearly express CD25 (Fig. 4A). At 4-8 dpi, the time when autocrine IL-2 signals are needed for CD4 T cell memory (47), substantial populations of CD11c^+^, MHC-II^+^ cells express CD25 in the lung, dLN and spleen (Fig 4B).

**Figure 4:**
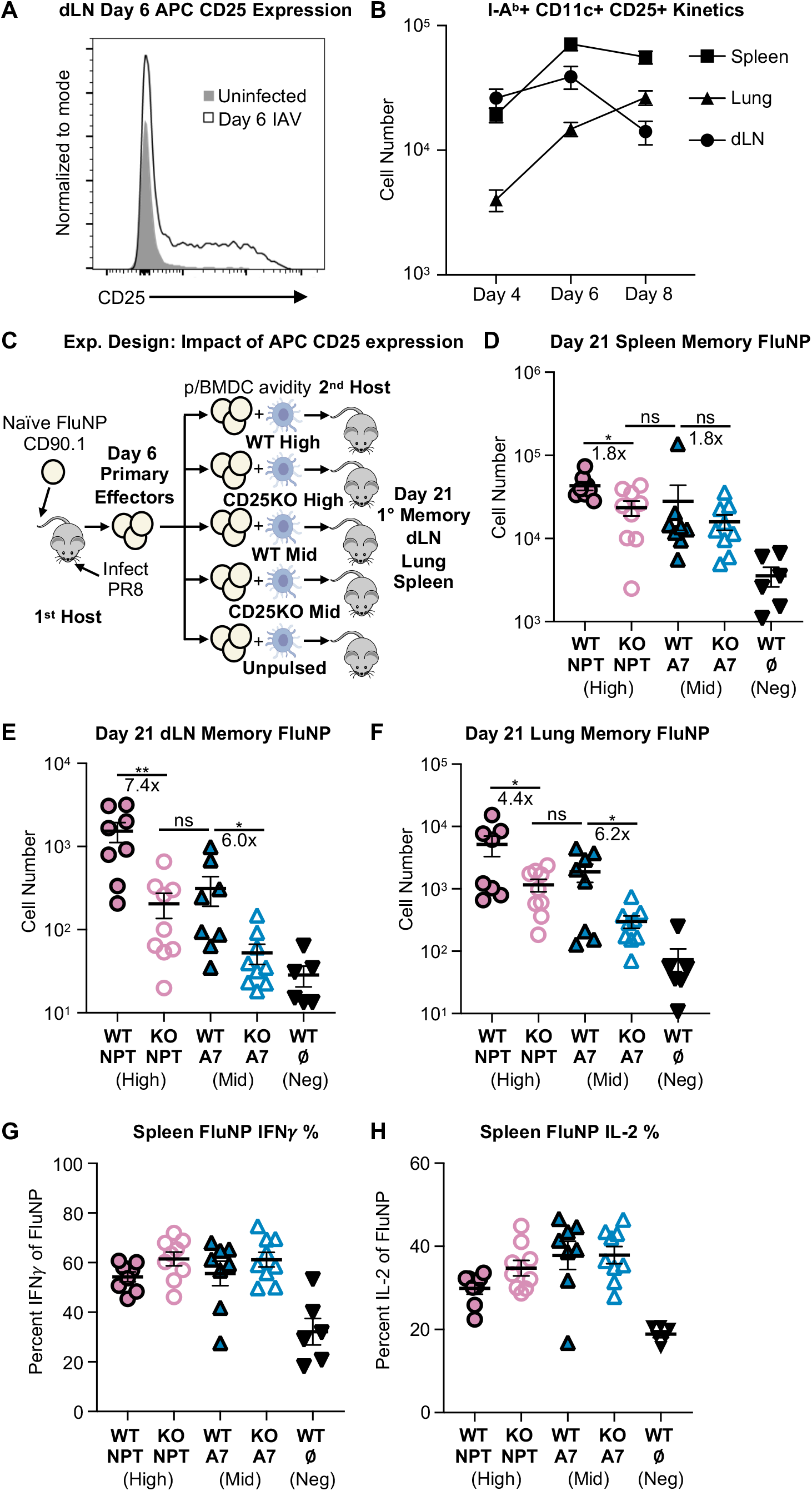
CD25 expression on APC induces more CD4 effectors to become memory. **(A-B)** CD25 Expression on APC *in vivo.* Mice were infected or not with PR8 influenza and sacrificed 4, 6 and 8 dpi. **(A)** CD25 expression was measured by FACS staining on I-A^b+^, CD11c^+^ cells in the dLN of infected (black line, no fill) or uninfected (gray) mice. **(B)** Kinetics of CD25^+^ I-A^b+^, CD11c^+^ cells in the dLN, lung, and spleen of infected mice at 4, 6 and 8 dpi. **(CH)** Impact of CD25 deletion in APC on memory generation. **(C)** Experimental Design. 6 dpi FluNP effectors were co-transferred with peptide-pulsed WT or CD25KO BMDC to uninfected 2^nd^ hosts. 2^nd^ hosts were sacrificed 15 dpt (21 dpi), and donor memory cells were analyzed by FACS. Number of memory FluNP cells determined by FACS analysis in **(D)** spleen, **(E)** dLN and **(F)** lung. **(G-H)** Spleen cells were re-stimulated with 10μM of NP_311-325_ for 6 hours *ex vivo* and percent expression of 21 dpi FluNP **(G)** IFNγ and **(H)** IL-2 determined by ICCS. Gated on live singlets, CD4^+^, CD90.1^+^, CD44^hi^. (A) Representative data, n = 10, two experiments. (B) Pooled data, n = 9-10, two experiments, mean +/- SEM. (D-H) Pooled data, n = 6-9, two experiments, mean +/- SEM. Statistical significance determined by two-tailed independent t test (*p<0.05, **p<0.01).

To analyze whether CD25 expression on APC plays a role in CD4 effector survival to memory, we generated 6d FluNP effectors and co-transferred them with WT or CD25KO activated BMDC pulsed with high and mid NP peptides (Fig. 4C). Both APC expressed high levels of CD11c, MHC-II, CD80 and CD86, but only WT BMDC expressed CD25 (Supp. Fig. 4A). We found that 6d effectors stimulated *in vitro* with WT vs. CD25KO peptide/APC produced equivalent amounts of IL-2 and IFNγ (Supp. Fig 4B-C) which indicates both activate FluNP equivalently. However, when we assessed *in vivo* generation of memory in the transfer model with WT or CD25KO APC (Fig. 4C), significantly fewer donor memory cells developed in spleen, dLN and lung when NPT high peptide was presented by CD25KO BMDC (Fig. 4D-F). With the middle avidity peptide there were significantly fewer memory cells generated with CD25KO BMDC in dLN and lung. The fraction of donor memory cells that produced IL-2, IFNγ and TNFα was equivalent (Fig 4G-H, Supp. Fig 4D). This suggests that transpresentation of IL-2 by APC serves to amplify the IL-2 signal to the CD4 effectors leading to generation of more memory cells.

### Peptide avidity at the effector checkpoint determines protection to influenza challenge

We asked if mice which received higher avidity peptide were better protected from rechallenge with IAV. We generated memory from FluNP effectors co-transferred with high or low Ag/APC (Fig. 5A). After 21 dpi, we challenged the mice with a sublethal dose of PR8 and analyzed their weight loss (Fig. 5B). The high avidity group recovered weight significantly faster than the low group, and both recovered more rapidly than the unpulsed group (Fig. 5B). When we challenged hosts with a higher, lethal dose of PR8 and analyzed survival, more animals in the two high groups (>80%) survived compared to their two low signaling counterparts (50-60%) and these were better protected than unpulsed APC (20%) (Fig. 5C). Thus, higher avidity peptide at the effector phase promoted a more protective memory FluNP population and the degree of protection increased with the size of the memory population.

**Figure 5:**
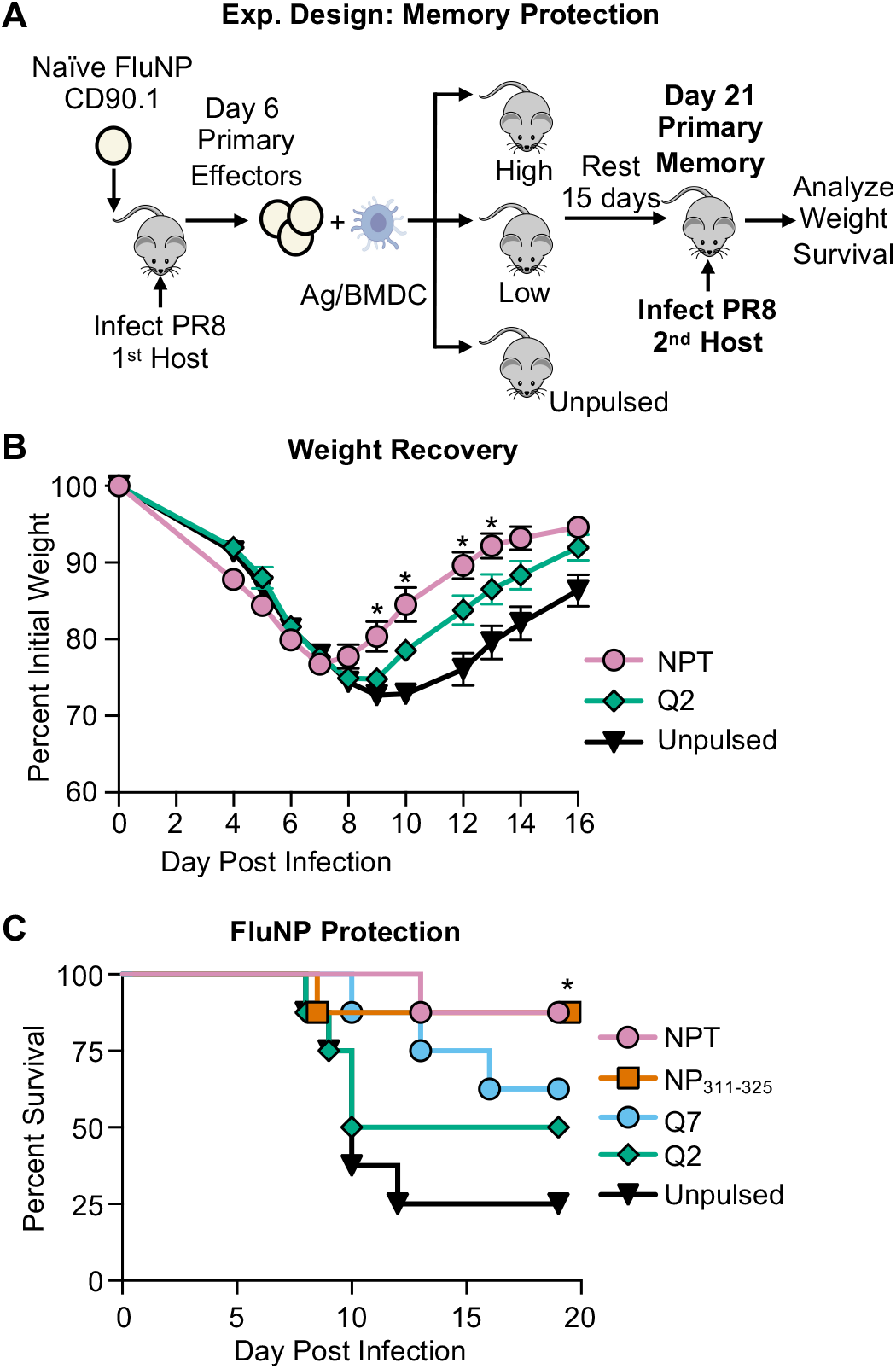
Increased peptide avidity at the effector checkpoint promotes a more protective population of memory cells. **(A)** Experimental design: FluNP effector cells (6 dpi) were co-transferred with peptide-pulsed APC into uninfected 2^nd^ hosts and rested for 15 d (21 dpi). At 21 dpi, 2^nd^ hosts were challenged with PR8. **(B)** Weight loss was determined following 2LD_50_ PR8 challenge. Statistical significance determined by two-tailed independent t test (*p<0.05). **(C)** Survival was measured following 4LD_50_ PR8 challenge. Statistical significance determined by log-rank (Mantel-Cox) test (*p<0.05). (B) Pooled data, n= 10-15, two experiments, mean +/- SEM. (C) Pooled data, n= 8, two experiments. (E-G) Pooled data, n= 8-9, two experiments, mean +/- SEM.

### Ag recognition by secondary effectors is required for memory generation and peptide avidity determines the size of response

Memory T cells have less stringent requirements than naïve for Ag dose and costimulatory interactions (48), and they become more protective secondary effectors (49). Memory T cells to influenza accumulate with age in humans due to multiple exposures, through both influenza infections and vaccinations (50–52) and thus may dominate responses. We asked whether secondary (2°) effectors generated from memory cells, also require Ag recognition to become secondary memory. We generated primary memory cells from 6d FluNP effectors with high peptide/APC in the 2^nd^ host to maintain the otherwise naïve state. At 21d, we infected 2°hosts and isolated 6d donor 2°effectors. We then co-transferred 2°6d FluNP effectors along with high, mid and low avidity peptide pulsed APC or unpulsed APC (Fig 6A) into 3^rd^ hosts and enumerated 2°memory FluNP after 21d in the spleen, dLN and lung (Fig. 6B-G). In each organ, memory generation was dependent on Ag recognition, with the high peptide NPT producing 14.4-fold more memory in spleen, 40-fold more in dLN and 34-fold more in lung compared to APC with no peptide Ag. As in the primary response, the number of memory cells was also reduced when lower affinity peptides were used for pulsing APC. Thus, CD4 memory cells, like naïve cells, need to recognize Ag again as secondary effectors, to form optimal secondary memory and peptide avidity again determines how many secondary memory cells are formed.

**Figure 6:**
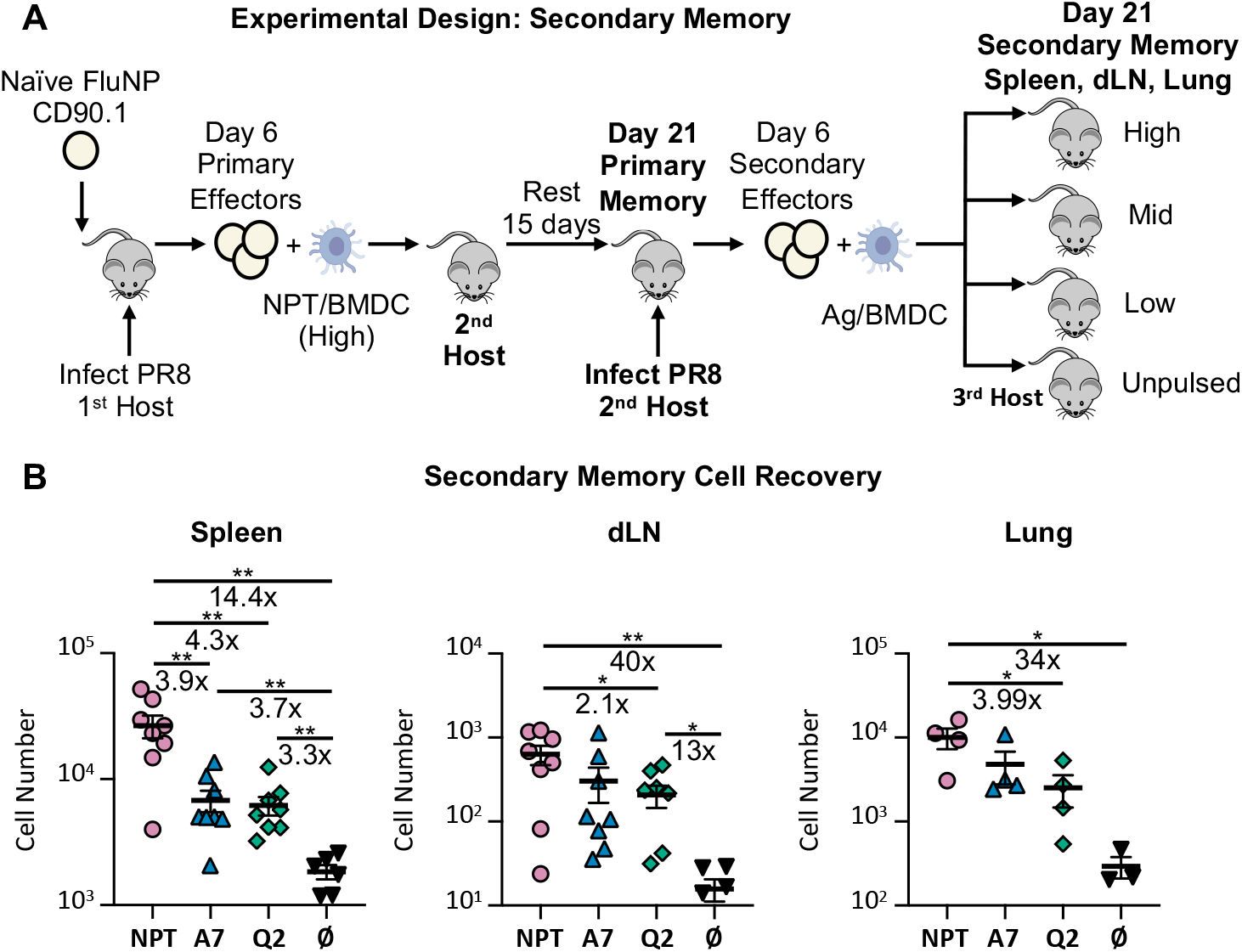
Peptide avidity during the secondary effector checkpoint regulates the size of the secondary memory population. **(A)** Experimental design. Primary 6 dpi FluNP effectors were generated and 1.5×10^6^ cotransferred to 2^nd^ hosts with 10^6^ NPT (high) peptide-pulsed BMDCs. At the memory stage, 21 dpi for the donor cells, 2^nd^ hosts were infected with PR8 influenza (0.3 LD_50_) and secondary 6 dpi effectors were isolated via CD90.1 MACS and co-transferred to uninfected 3^rd^ hosts. Fifteen days later 3^rd^ hosts were sacrificed, and donor secondary memory cells were analyzed by FACS. **(B)** Number of secondary memory donor FluNP cells determined FACS analysis in the spleen, dLN and lung. Spleen and dLN pooled data, n = 6-8, two experiments, mean +/- SEM. Lung representative data n = 6-8, two experiments, mean +/- SEM. Statistical significance determined by two-tailed independent t test (*p<0.05, **p<0.01).

## Discussion

We examined the impact of peptide avidity for the FluNP TCR, delivered at the CD4 effector checkpoint (23, 24), on memory formation. We recovered the greatest number of CD4 memory cells after 6d effecters were stimulated with the highest avidity peptide, with 200-fold more in the lung compared to no peptide, and 20-50-fold more compared to low avidity peptide. Higher avidity drove higher levels of autocrine IL-2 production, which promoted greater effector survival and donor cell recovery in the late effector phase, laying out the mechanisms likely to be responsible. The importance of autocrine IL-2 in memory formation was emphasized by the fact that optimum levels of memory required that activated APC express IL-2Rα during the cognate interaction with CD4 effectors. Higher avidity also led to increased protection from lethal rechallenge. 2°CD4 effector cells also required Ag recognition to form secondary memory and were favored by high avidity peptide. To increase CD4 T cell memory, we suggest vaccine strategies must supply a second round of high avidity Ag and PR signals shortly after initial immunization during the effector T cell response.

Previous studies varied Ag avidity at the initiation of the immune response and found higher levels and avidity could led to greater effector and memory cell number (53–56). Extending the period of Ag presentation also promoted enhanced CD4 and CD8 memory formation (2–4, 23, 24, 57, 58). The pathways responsible were not determined and no studies restricted Ag to the effector checkpoint, that we know is strictly required for memory generation (23, 24, 47). Our study is the first to analyze the effect of a broad range of affinities during the effector checkpoint on memory. Our results indicate that at the effector checkpoint 6dpi after initial infection, peptide/MHC-II avidity for the TCR on the CD4 effectors during cognate interaction determines the magnitude of memory by regulating the level of effector production of autocrine IL-2. In turn, this determines how many effectors survive over the next few days and progress to memory. In the polyclonal situation, this likely selects for CD4 memory cells which are of the highest affinity for the pathogen Ag they recognize.

In our studies, the impact of peptide avidity on memory was consistently more dramatic in lung and dLN, compared to the spleen. One likely explanation for this is that higher avidity peptide also promotes effector migration from the spleen to the lung, so the impact is less apparent in the spleen. We and others have shown that checkpoint Ag recognition and strong Th1 skewing promote greater expression of CXCR3 by effector cells promoting their trafficking to tissue sites (23, 35, 59, 60). The results are particularly striking here, because in the 2^nd^ host there is no infection or inflammation in the lung to attract effectors. Thus, we suggest peptide avidity at the effector checkpoint also regulates migration to the tissues.

We found previously that Ag presentation to effector CD4 T cells at the effector checkpoint induces autocrine IL-2 production, which prevents their default apoptosis and thus supports formation of memory cells (16, 17). Here, we show higher peptide avidity does not drive greater division of 6d effectors, but does proportionally increase their level of IL-2 production and leads to a dramatic increase in short term effector survival and recovery, such that the number of effectors at 9d (Fig. 3D) is proportional to the size of the memory population at 21d (Fig. 2E). In each organ, there were more than 100-fold higher donor effector cells recovered with the highest peptide vs. no peptide, with very few donor cells in the unpulsed groups (spleen ~10^3^, dLN ~500 and lung ~50). At this timepoint even the low avidity peptide results in 20-40-fold more effector cells than no peptide (Fig. 3D). The increase in IL-2 production by effectors at 6d with avidity and dose of Ag (Fig. 3), clearly links the effect of peptide avidity to IL-2 rescue of effectors and thus increased memory formation.

However, CD25 was either transiently expressed or not expressed on most CD4 effectors, raising the possibility that optimum autocrine IL-2-mediated survival of effectors might be regulated by something in addition to IL-2 binding to IL-2R complex on the CD4 T cell. Another survival cytokine, IL-15, which shares IL-2Rβ/γ with IL-2, is presented *in trans* when bound to IL-15Rα on APC while signaling through IL-2Rβ/γ on the T cell (41–43) and previous studies reported that activated APC can express CD25 in both mice and humans (44, 61). We found the activated BMDC we use as APC, as well as a subset of activated APC in mice infected with IAV 4-8 d earlier, express high levels of CD25 (Fig 4). We tested CD25 deficient APC and found they drove less memory formation compared to WT APC *in vivo* (Fig. 4). This suggests that Ag/APC transpresent autocrine IL-2 to the interacting CD4 T cells increasing IL-2 availability at the time CD4 effectors do not express CD25, driving greater effector survival and memory formation. Thus, we suggest infection provided pathogen recognition (PR) signals that activate APC, enhancing their CD25 expression, as well as MHC-II and costimulatory ligands, so they efficiently present both peptide Ag and CD4 effector-produced IL-2. We suggest this mechanism evolved to require that infection provide high avidity Ag at a high dose, and APC activation by PR signals, to promote optimum effector transition to memory, especially in the lung, the site of infection. This mechanism would restrict memory generation to situations where virus is still replicating and providing PR signals at the effector stage. Thus, effective vaccine strategies likely need to provide high dose, high avidity Ag and PR signals again during the T cell effector phase.

In adult humans, with a long history of exposure to influenza viruses, many responses likely stem from existing memory cells (50–52). We find that efficient generation of 2°memory also requires effector checkpoint Ag recognition and is increased by high avidity interactions (Fig. 6). This contrasts with observations that memory responses in general are less dependent on Ag dose and costimulation (49). We suggest that effector functions of memory cells are more easily achieved, but when forming new 2°memory, stringent regulation is again in place to avoid unnecessary memory cells and select only those with high affinity for persistent Ag in the context of infection.

The impressive impact of Ag avidity for TCR in determining the size of the memory pool from both primary (Fig. 2) and secondary (Fig. 6) CD4 effectors indicates that both a high dose of available peptides and CD4 effectors bearing TCR with high affinity for some of those peptides are strictly required for robust memory generation. This implies immunization with a wide range of viral proteins, including those with known immunodominant CD4 epitopes, will be more effective in inducing memory then single proteins. Because of the heterogeneity of human class-II, only a fraction of total potential viral epitopes will be immunodominant in each individual, which also argues for vaccines expressing a breadth of proteins. A broader repertoire of high affinity memory CD4 T cells also should lower the likelihood that escape variants, which arise by random mutations and selection, will have the opportunity to develop and escape a broad CD4 T cell response so they can be passed on, since this would need require multiple mutations. Many of the immunodominant T cell epitopes, such as the FluNP NP_311_ used here, are in core proteins of viruses, not in the viral surface proteins that B cells recognize, thus it follows that vaccines should include both core and surface protein epitopes, to elicit both T and B cell immune memory including heterosubtypic determinants not likely to be selected by Ab to external surface proteins.

Since more memory is generated when CD25 is expressed by activated APC, our studies re-affirm that vaccines need to provide continuing PR signals that directly activate APC. Our results predict that in polyclonal responses, the requirement for high avidity of TCR for Ag/APC will serve to select a spectrum of memory cells of higher affinity than the effector population. This should result in more effective protection in the real-life polyclonal situation. An advantage of vaccines is that they can supply these signals in a less dangerous context than infection. Additional studies are needed to support these later implications in detail, but we suggest that the overall purpose of the requirement for a high avidity interaction at the effector stage is to provide more memory cells with higher affinity for the infecting virus, while not allowing memory when virus does not persist at high levels into the effector stage, or there is no replicating infectious entity.

---

**Supplementary Figure 1. Comparison of FluNP vs. CD4 host response: cytokines, phenotype and memory subsets.**

Naïve FluNP.Thy1.1^+/-^ cells were transferred to B6 hosts, that were then infected with PR8. **(A)** Kinetics of FluNP Response (SLO) Lung draining lymph node (dLN) and spleen were collected at 4, 6, 8, 12, 21 and 63 dpi and the number of donor FluNP cells determined in each mouse by FACS analysis. **(B-E)** Representative FACS plots for effector subset analysis (8 dpi) gated on lung donor FluNP and responding host CD4^+^ cells: **(B)** T-bet^+^ CD44^+^, **(C)** NKG2A/C/E^+^ CD44^+^, **(D)** FoxP3^+^ CD44^+^ and **(E)** cytokines (IFNγ^+^, TNFα^+^, IL-2^+^, IL-17^+^). **(F-I)** Representative FACS plots for effector subset analysis (8 dpi) gated on spleen donor FluNP and responding host CD4^+^ cells: **(F)** T-bet^+^ CD44^+^, **(G)** CXCR5 BCL-6^+^, **(H)** FoxP3^+^ CD44^+^ and **(I)** cytokines (IFNγ^+^, TNFα^+^, IL-2^+^, IL-17^+^). **(J-M)** Phenotyping of memory subsets (21 dpi). Representative FACS plots gated on donor FluNP and responding host CD4^+^ cells: **(J)** lung CD69^+^ CD44^+^ and **(K)** spleen CD44^+^ CD127^+^. **(L)** Percent CD127 of CD44^hi^ donor FluNP and responding host CD4^+^, CD44^hi^ in dLN, lung and spleen. **(M)** Representative CD62L FACS plots gated on day 21 spleen donor FluNP, CD44^hi^, CD127^+^ and host CD4^+^, CD44^hi^, CD127^+^. (A) Day 4, 6, 8, 21, 63 Pooled data, n = 9-10, two experiments. Day 12 one experiment n = 5. Mean +/- SEM. (B-I) Representative data, n = 11, two experiments. (J, K, M) Representative data, n = 7, two experiments. (L) Pooled data, n = 7, two experiments. Statistical significance determined by two-tailed independent t test (***p<0.001).

**Supplementary Figure 2. Related to Figure 2. Memory FluNP gating, phenotyping and cytokine production.**

**(A-H)** FluNP 6d effector cells were generated as in Figure 2 and 1.5×10^6^ 6d FluNP effectors were co-transferred with Ag/APC into uninfected 2^nd^ hosts. Fifteen days later (21 dpi) 2^nd^ hosts were sacrificed, and the donor FluNP memory cells were analyzed by FACS. **(A)** Representative FACS plots showing donor memory cells by CD44 and CD90.1 expression in the spleen at 21 dpi (Fig. 2D cont). **(B)** Representative FACS plots of CD44 and CD127 expression of donor FluNP. Gated on live singlets, CD4^+^, CD90.1^+^ cells. **(C-E)** Percent of donor FluNP expressing CD127 on day at 21 dpi in **(C)** spleen **(D)** dLN and **(E)** lung. **(F-H)** Representative histograms (left), percent expression (middle) and normalized median fluorescence intensity (nMFI) (right) of ICCS (**F**) IFNγ^+^ **(G)** TNFα^+^ **(H)** IL-2^+^. Gated on live singlets, CD4^+^, CD90.1^+^, CD44^hi^ cells from spleen 21 dpi. Statistical significance determined by two-tailed independent t test (*p<0.05, **p<0.01, ***p<0.001, ****p<0.0001). (A-B) Representative data, n = 8-15, four experiments. (C-E) Pooled data, n = 8-15, four experiments, mean +/- SEM. (F-H) Representative data (histograms) and pooled data, n = 8-15, four experiments, mean +/- SEM.

**Supplementary Figure 3. Related to Figure 3. Donor cell survival and proliferation and IL-2R subunit expression 3 days post transfer, day 6 cytokine production.**

**(A-C)** <u>Division in spleen</u>. FluNP effectors (6 dpi) were labeled with CTV and co-transferred with 10^6^ peptide-pulsed BMDCs into 2^nd^ hosts. 2^nd^ hosts were sacrificed 3 dpt and donor FluNP CTV expression was analyzed. **(A)** Representative histogram of donor FluNP CTV dilution 3 dpt in spleen. Gates 1-5 are equally sized to undivided. **(B-C)** Percent of donor FluNP cells 3 dpt within gates shown in (A): **(B)** spleen, **(C)** dLN and lung. **(D)** <u>Survival in spleen</u>. Representative FACS plots of donor FluNP survival by live/dead yellow and FLICA caspase 3/7 staining in spleen 3 dpt. Fraction of live donor FluNP by FACS in the **(E)** dLN and **(F)** lung. **(G)** Expression of IL-2R chains by CD44^hi^ FluNP donor cells in spleen, dLN and lung 3 dpt. **(H-J)** FluNP 6 dpi effectors were restimulated for 6 hr with indicated NP peptides at 10μM and **(H)** TNFα and **(I)** IFNγ expression determined by FACS. **(J)** FluNP effectors were co-cultured with NP peptide-pulsed APC for 48 hr and IFNγ production in the supernatant was determined by ELISA. Statistics: (A, D, G-I) Representative data, n = 10, two experiments. (B-C) Pooled data, n = 10, two experiments, mean +/- SEM. (K, M) Pooled data, n = 10, two experiments, mean +/- SEM. (L) Pooled data, n = 3-6, two experiments, mean +/- SEM. Statistical significance determined by two-tailed independent t test (*p<0.05, **p<0.01, ***p<0.001).

**Supplementary Figure 4. Related to Figure 4. Characterization of WT and CD25KO APC, and their impact on induction of FluNP effector cytokine induction.**

**(A)** Histograms of CD11c, I-A^b^, CD86, CD80 and CD25 expression on BMDC prepared from WT and CD25KO mice. **(B-C)** FluNP 6d effectors were stimulated with WT or CD25KO APC pulsed with NPT (high) or A7 (mid) NP peptide for 6 hr and production of **(B)** IL-2 and **(C)** IFNγ determined by ELISA. **(D)** Spleen cells (Fig. 4C) were re-stimulated with 10μM of NP_311-325_ for 6 hours *ex vivo* and percent expression of 21 dpi FluNP TNFα determined by FACS. Gated on live singlets, CD4^+^, CD90.1^+^, CD44^hi^. (A) Representative data, n = 2, two experiments. (B-C) Pooled data, n =2-3, three experiments, mean +/- SEM. (D) Pooled data, n = 6-9, two experiments, mean +/- SEM. (E-G) Pooled data, n = 8-9, two experiments, mean +/- SEM.

## Notes

### Competing Interest Statement

The authors have declared no competing interest.

## References

1. McKinstry, K. K., T. M. Strutt, Y. Kuang, D. M. Brown, S. Sell, R. W. Dutton, and S. L. Swain. 2012. Memory CD4+ T cells protect against influenza through multiple synergizing mechanisms. J Clin Invest 122: 2847–2856.

2. Obst, R., H.-M. van Santen, D. Mathis, and C. Benoist. 2005. Antigen persistence is required throughout the expansion phase of a CD4+ T cell response. J Exp Med 201: 1555–1565.

3. Prlic, M., G. Hernandez-Hoyos, and M. J. Bevan. 2006. Duration of the initial TCR stimulus controls the magnitude but not functionality of the CD8+ T cell response. J Exp Med 203: 2135–2143.

4. Rabenstein, H., A. C. Behrendt, J. W. Ellwart, R. Naumann, M. Horsch, J. Beckers, and R. Obst. 2014. Differential kinetics of antigen dependency of CD4+ and CD8+ T cells. J Immunol 192: 3507–3517.

5. Corse, E., R. A. Gottschalk, and J. P. Allison. 2011. Strength of TCR-Peptide/MHC Interactions and In Vivo T Cell Responses. J Immunol 186: 5039–5045.

6. Hosken, B. N. A., K. Shibuya, A. W. Heath, K. M. Murphy, and A. O. Garra. 1995. The effect of antigen dose on CD4+ T helper cell phenotype development in a T cell receptor-alpha beta-transgenic model. J Exp Med 182: 1579–1584.

7. Brogdon, J. L., D. Leitenberg, and K. Bottomly. 2002. The Potency of TCR Signaling Differentially Regulates NFATc/p Activity and Early IL-4 Transcription in Naive CD4+ T Cells. J Immunol 168: 3825–3832.

8. Constant, B. S., C. Pfeiffer, A. Woodard, T. Pasqualini, and K. Bottomly. 1995. Extent of T cell receptor ligation can determine the functional differentiation of naive CD4+ T cells. J Exp Med 182: 1591–1596.

9. Tubo, N. J., A. J. Pagán, J. J. Taylor, R. W. Nelson, J. L. Linehan, J. M. Ertelt, E. S. Huseby, S. S. Way, and M. K. Jenkins. 2013. Single Naive CD4+ T Cells from a Diverse Repertoire Produce Different Effector Cell Types during Infection. Cell 153: 785–796.

10. Fazilleau, N., L. J. McHeyzer-Williams, H. Rosen, and M. G. McHeyzer-Williams. 2009. The function of follicular helper T cells is regulated by the strength of T cell antigen receptor binding. Nat Immunol 10: 375.

11. Keck, S., M. Schmaler, S. Ganter, L. Wyss, S. Oberle, E. S. Huseby, D. Zehn, and C. G. King. 2014. Antigen affinity and antigen dose exert distinct influences on CD4 T-cell differentiation. Proc Natl Acad Sci USA 111: 14852–14857.

12. Kotov, D. I., J. S. Mitchell, T. Pengo, C. Ruedl, S. S. Way, R. A. Langlois, B. T. Fife, and M. K. Jenkins. 2019. TCR Affinity Biases Th Cell Differentiation by Regulating CD25, Eef1e1, and Gbp2. J Immunol 202: 2535–2545.

13. Snook, J. P., C. Kim, and M. A. Williams. 2018. TCR signal strength controls the differentiation of CD4+ effector and memory T cells. Sci Immunol 3: 1–13.

14. Busch, D. H., I. Pilip, and E. G. Pamer. 1998. Evolution of a complex T cell receptor repertoire during primary and recall bacterial infection. J Exp Med 188: 61–70.

15. Crawford, F., H. Kozono, J. White, P. Marrack, and J. Kappler. 1998. Detection of antigen-specific T cells with multivalent soluble class II MHC covalent peptide complexes. Immunity 8: 675–682.

16. Busch, D. H., and E. G. Pamer. 1999. T cell affinity maturation by selective expansion during infection. J Exp Med 189: 701–709.

17. Savage, P. A., J. J. Boniface, and M. M. Davis. 1999. A kinetic basis for T cell receptor repertoire selection during an immune response. Immunity 10: 485–492.

18. Sabatino, J. J., Jr., J. Huang, C. Zhu, and B. D. Evavold. 2011. High prevalence of low affinity peptide-MHC II tetramer-negative effectors during polyclonal CD4+ T cell responses. J Exp Med 208: 81–90.

19. Malherbe, L., C. Hausl, L. Teyton, and M. G. McHeyzer-Williams. 2004. Clonal selection of helper T cells is determined by an affinity threshold with no further skewing of TCR binding properties. Immunity 21: 669–679.

20. Corse, E., R. A. Gottschalk, M. Krogsgaard, and J. P. Allison. 2010. Attenuated T cell responses to a high-potency ligand in vivo. PLoS Biol 8.

21. Xia, J., Y. Kuang, J. Liang, M. Jones, and S. L. Swain. 2020. Influenza Vaccine-Induced CD4 Effectors Require Antigen Recognition at an Effector Checkpoint to Generate CD4 Lung Memory and Antibody Production. J Immunol 205: 2077–2090.

22. Powell, T. J., T. Strutt, J. Reome, J. A. Hollenbaugh, A. D. Roberts, D. L. Woodland, S. L. Swain, and R. W. Dutton. 2007. Priming with cold-adapted influenza A does not prevent infection but elicits long-lived protection against supralethal challenge with heterosubtypic virus. J Immunol 178: 1030–1038.

23. Bautista, B. L., P. Devarajan, K. K. McKinstry, T. M. Strutt, A. M. Vong, M. C. Jones, Y. Kuang, D. Mott, and S. L. Swain. 2016. Short-Lived Antigen Recognition but Not Viral Infection at a Defined Checkpoint Programs Effector CD4 T Cells To Become Protective Memory. J Immunol 197: 3936–3949.

24. McKinstry, K. K., T. M. Strutt, B. Bautista, W. Zhang, Y. Kuang, A. M. Cooper, and S. L. Swain. 2014. Effector CD4 T-cell transition to memory requires late cognate interactions that induce autocrine IL-2. Nat Commun 5: 5377.

25. Zhang, B. X., L. Giangreco, H. E. Broome, C. M. Dargan, and S. L. Swain. 1995. Control of CD4 effector fate: transforming growth factor beta 1 and interleukin 2 synergize to prevent apoptosis and promote effector expansion. J Exp Med 182: 699–709.

26. Moran, A. E., K. L. Holzapfel, Y. Xing, N. R. Cunningham, J. S. Maltzman, J. Punt, and K. A. Hogquist. 2011. T cell receptor signal strength in Treg and iNKT cell development demonstrated by a novel fluorescent reporter mouse. J Exp Med 208: 1279–1289.

27. Stadinski, B. D., P. Trenh, R. L. Smith, B. Bautista, P. G. Huseby, G. Li, L. J. Stern, and E. S. Huseby. 2011. A role for differential variable gene pairing in creating T cell receptors specific for unique major histocompatibility ligands. Immunity 35: 694–704.

28. Huseby, E. S., B. Sather, P. G. Huseby, and J. Goverman. 2001. Age-dependent T cell tolerance and autoimmunity to myelin basic protein. Immunity 14: 471–481.

29. Zhumabekov, T., P. Corbella, M. Tolaini, and D. Kioussis. 1995. Improved version of a human CD2 minigene based vector for T cell-specific expression in transgenic mice. Journal of Immunological Methods 185: 133–140.

30. Brahmakshatriya, V., Y. Kuang, P. Devarajan, J. Xia, W. Zhang, A. M. Vong, and S. L. Swain. 2017. IL-6 Production by TLR-Activated APC Broadly Enhances Aged Cognate CD4 Helper and B Cell Antibody Responses In Vivo. J Immunol 198: 2819–2833.

31. Marshall, N. B., A. M. Vong, P. Devarajan, M. D. Brauner, Y. Kuang, R. Nayar, E. A. Schutten, C. H. Castonguay, L. J. Berg, S. L. Nutt, and S. L. Swain. 2017. NKG2C/E Marks the Unique Cytotoxic CD4 T Cell Subset, ThCTL, Generated by Influenza Infection. J Immunol 198: 1142–1155.

32. Devarajan, P., A. M. Vong, C. H. Castonguay, O. Kugler-Umana, B. L. Bautista, M. C. Jones, K. A. Kelly, J. Xia, and S. L. Swain. 2022. Strong influenza-induced TFH generation requires CD4 effectors to recognize antigen locally and receive signals from continuing infection. Proc Natl Acad Sci USA 119.

33. McKinstry, K. K., S. Golech, W. H. Lee, G. Huston, N. P. Weng, and S. L. Swain. 2007. Rapid default transition of CD4 T cell effectors to functional memory cells. J Exp Med 204: 2199–2211.

34. McKinstry, K. K., T. M. Strutt, and S. L. Swain. 2010. Regulation of CD4+ T-cell contraction during pathogen challenge. Immunological Reviews 236: 110–124.

35. Dhume, K., C. M. Finn, T. M. Strutt, S. Sell, and K. K. McKinstry. 2019. T-bet optimizes CD4 T-cell responses against influenza through CXCR3-dependent lung trafficking but not functional programming. Mucosal Immunol 12: 1220–1230.

36. Nanaware, P. P., M. M. Jurewicz, C. C. Clement, L. Lu, L. Santambrogio, and L. J. Stern. 2021. Distinguishing Signal From Noise in Immunopeptidome Studies of Limiting-Abundance Biological Samples: Peptides Presented by I-A(b) in C57BL/6 Mouse Thymus. Front Immunol 12: 658601.

37. Itoh, Y., K. Kajino, K. Ogasawara, A. Takahashi, K. Namba, I. Negishi, N. Matsuki, K. Iwabuchi, M. Kakinuma, R. A. Good, and K. Onoé. 1997. Interaction of pigeon cytochrome c-(43-58) peptide analogs with either T cell antigen receptor or I-Ab molecule. Proc Natl Acad Sci U S A 94: 12047–12052.

38. Zhu, Y., A. Y. Rudensky, A. L. Corper, L. Teyton, and I. A. Wilson. 2003. Crystal structure of MHC class II I-Ab in complex with a human CLIP peptide: prediction of an I-Ab peptide-binding motif. J Mol Biol 326: 1157–1174.

39. Li, J., G. Huston, and S. L. Swain. 2003. IL-7 promotes the transition of CD4 effectors to persistent memory cells. J Exp Med 198: 1807–1815.

40. Bradly, L. M., D. D. Duncan, S. Tonkonogy, and S. L. Swain. 1991. Characterization of antigen-specific CD4+ effector T cells in vivo: immunization results in a transient population of MEL-14-, CD45RB-helper cells that secretes interleukin 2 (IL-2), IL-3, IL-4, and interferon gamma. J Exp Med 174: 547–559.

41. Lodolce, J. P., P. R. Burkett, D. L. Boone, M. Chien, and A. Ma. 2001. T cell-independent interleukin 15Rα signals are required for bystander proliferation. J Exp Med 194: 1187–1193.

42. Dubois, S., J. Mariner, T. A. Waldmann, and Y. Tagaya. 2002. IL-15Ra Recycles and Presents IL-15 In trans to Neighboring Cells. Immunity 17: 537–547.

43. Stonier, S. W., and K. S. Schluns. 2010. Trans-presentation: a novel mechanism regulating IL-15 delivery and responses. Immunol Lett 127: 85–92.

44. Wuest, S. C., J. H. Edwan, J. F. Martin, S. Han, J. S. Perry, C. M. Cartagena, E. Matsuura, D. Maric, T. A. Waldmann, and B. Bielekova. 2011. A role for interleukin-2 transpresentation in dendritic cell-mediated T cell activation in humans, as revealed by daclizumab therapy. Nat Med 17: 604–609.

45. Eicher, D. M., and T. A. Waldmann. 1998. IL-2Rα on one cell can present IL-2 to IL-2Rβ/γc on another cell to augment IL-2 signaling. J Immunol 161: 5430–5437.

46. Kim, M., T. J. Kim, H. M. Kim, J. Doh, and K. M. Lee. 2017. Multi-cellular natural killer (NK) cell clusters enhance NK cell activation through localizing IL-2 within the cluster. Sci Rep 7: 40623.

47. Swain, S. L., M. C. Jones, P. Devarajan, J. Xia, R. W. Dutton, T. M. Strutt, and K. K. McKinstry. 2021. Durable CD4 T-Cell Memory Generation Depends on Persistence of High Levels of Infection at an Effector Checkpoint that Determines Multiple Fates. Cold Spring Harb Perspect Biol 13.

48. Dubey, C., M. Croft, and S. L. Swain. 1995. Costimulatory requirements of naive CD4+ T cells. ICAM-1 or B7-1 can costimulate naive CD4 T cell activation but both are required for optimum response. J Immunol 155: 45–57.

49. Strutt, T. M., K. K. McKinstry, Y. Kuang, L. M. Bradley, and S. L. Swain. 2012. Memory CD4+ T-cell-mediated protection depends on secondary effectors that are distinct from and superior to primary effectors. Proc Natl Acad Sci USA 109: E2551–2560.

50. Beura, L. K., S. E. Hamilton, K. Bi, J. M. Schenkel, O. A. Odumade, K. A. Casey, E. A. Thompson, K. A. Fraser, P. C. Rosato, A. Filali-Mouhim, R. P. Sekaly, M. K. Jenkins, V. Vezys, W. N. Haining, S. C. Jameson, and D. Masopust. 2016. Normalizing the environment recapitulates adult human immune traits in laboratory mice. Nature 532: 512–516.

51. Reese, T. A., K. Bi, A. Kambal, A. Filali-Mouhim, L. K. Beura, M. C. Burger, B. Pulendran, R. P. Sekaly, S. C. Jameson, D. Masopust, W. N. Haining, and H. W. Virgin. 2016. Sequential Infection with Common Pathogens Promotes Human-like Immune Gene Expression and Altered Vaccine Response. Cell Host Microbe 19: 713–719.

52. Herati, R. S., A. Muselman, L. Vella, B. Bengsch, K. Parkhouse, D. Del Alcazar, J. Kotzin, S. A. Doyle, P. Tebas, S. E. Hensley, L. F. Su, K. E. Schmader, and E. J. Wherry. 2017. Successive annual influenza vaccination induces a recurrent oligoclonotypic memory response in circulating T follicular helper cells. Sci Immunol 2.

53. Vanguri, V., C. C. Govern, R. Smith, and E. S. Huseby. 2013. Viral antigen density and confinement time regulate the reactivity pattern of CD4 T-cell responses to vaccinia virus infection. Proc Natl Acad Sci USA 110: 288–293.

54. Kim, C., T. Wilson, K. F. Fischer, and M. A. Williams. 2013. Sustained interactions between T cell receptors and antigens promote the differentiation of CD4(+) memory T cells. Immunity 39: 508–520.

55. Cho, Y. L., M. Flossdorf, L. Kretschmer, T. Hofer, D. H. Busch, and V. R. Buchholz. 2017. TCR Signal Quality Modulates Fate Decisions of Single CD4(+) T Cells in a Probabilistic Manner. Cell Rep 20: 806–818.

56. Williams, M. A., E. V. Ravkov, and M. J. Bevan. 2008. Rapid culling of the CD4+ T cell repertoire in the transition from effector to memory. Immunity 28: 533–545.

57. León, B., A. Ballesteros-Tato, T. D. Randall, and F. E. Lund. 2014. Prolonged antigen presentation by immune complex-binding dendritic cells programs the proliferative capacity of memory CD8 T cells. J Exp Med 211: 1637–1655.

58. Ballesteros-Tato, A., B. Leon, B. O. Lee, F. E. Lund, and T. D. Randall. 2014. Epitope-specific regulation of memory programming by differential duration of antigen presentation to influenza-specific CD8(+) T cells. Immunity 41: 127–140.

59. Kohlmeier, J. E., T. Cookenham, S. C. Miller, A. D. Roberts, J. P. Christensen, A. R. Thomsen, and D. L. Woodland. 2009. CXCR3 directs antigen-specific effector CD4+ T cell migration to the lung during parainfluenza virus infection. J Immunol 183: 4378–4384.

60. Groom, J. R., and A. D. Luster. 2011. CXCR3 ligands: redundant, collaborative and antagonistic functions. Immunol Cell Biol 89: 207–215.

61. Li, J., E. Lu, T. Yi, and J. G. Cyster. 2016. EBI2 augments Tfh cell fate by promoting interaction with IL-2-quenching dendritic cells. Nature 533: 110–114.

