## Supplemental Figures for "Peptide Avidity for TCR on CD4 Effectors Determines the Extent of Memory Generation"

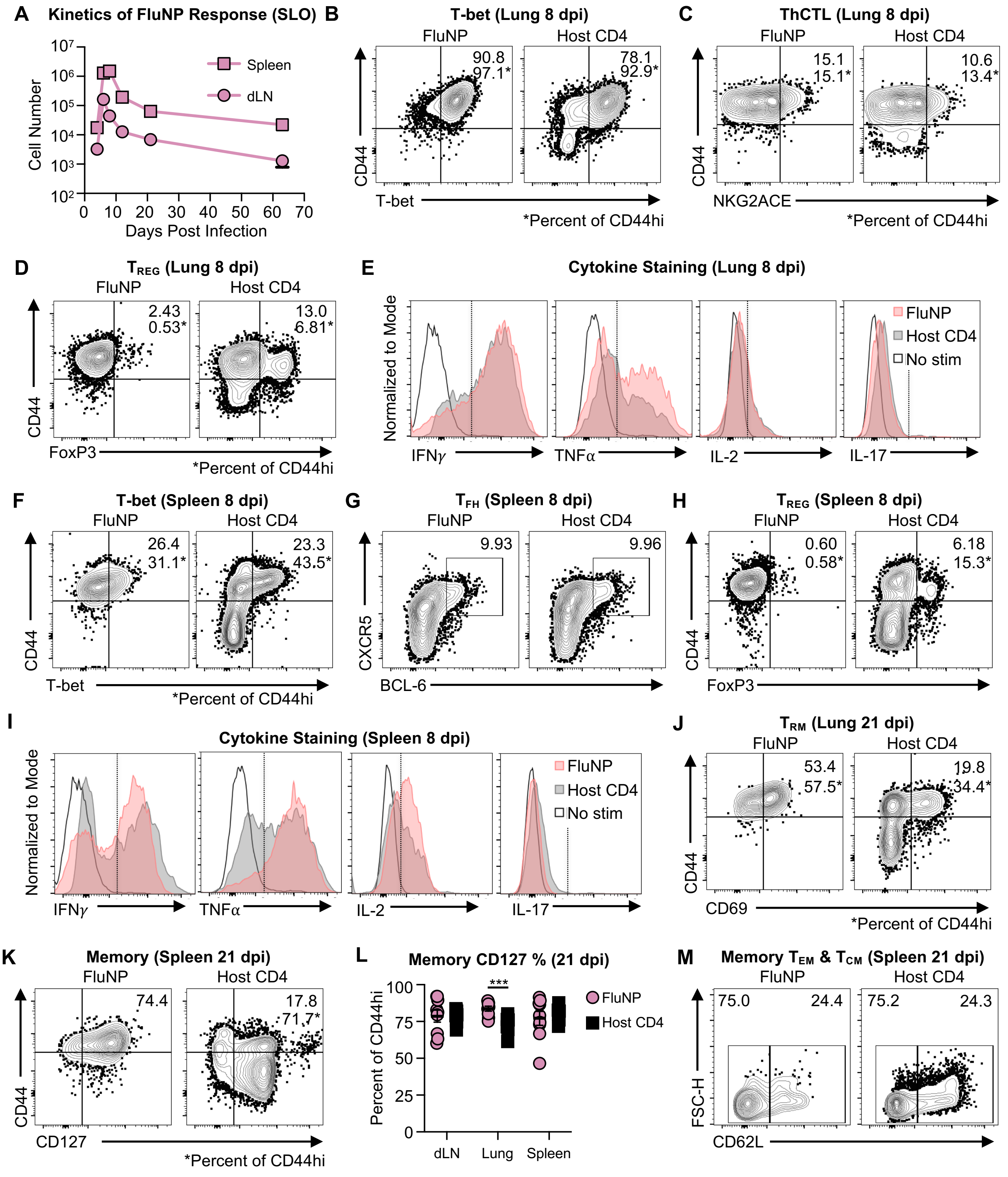


**Supplementary Figure 1. Related to Figure 1. Comparison of FluNP vs. CD4 host response: cytokines, phenotype and memory subsets**

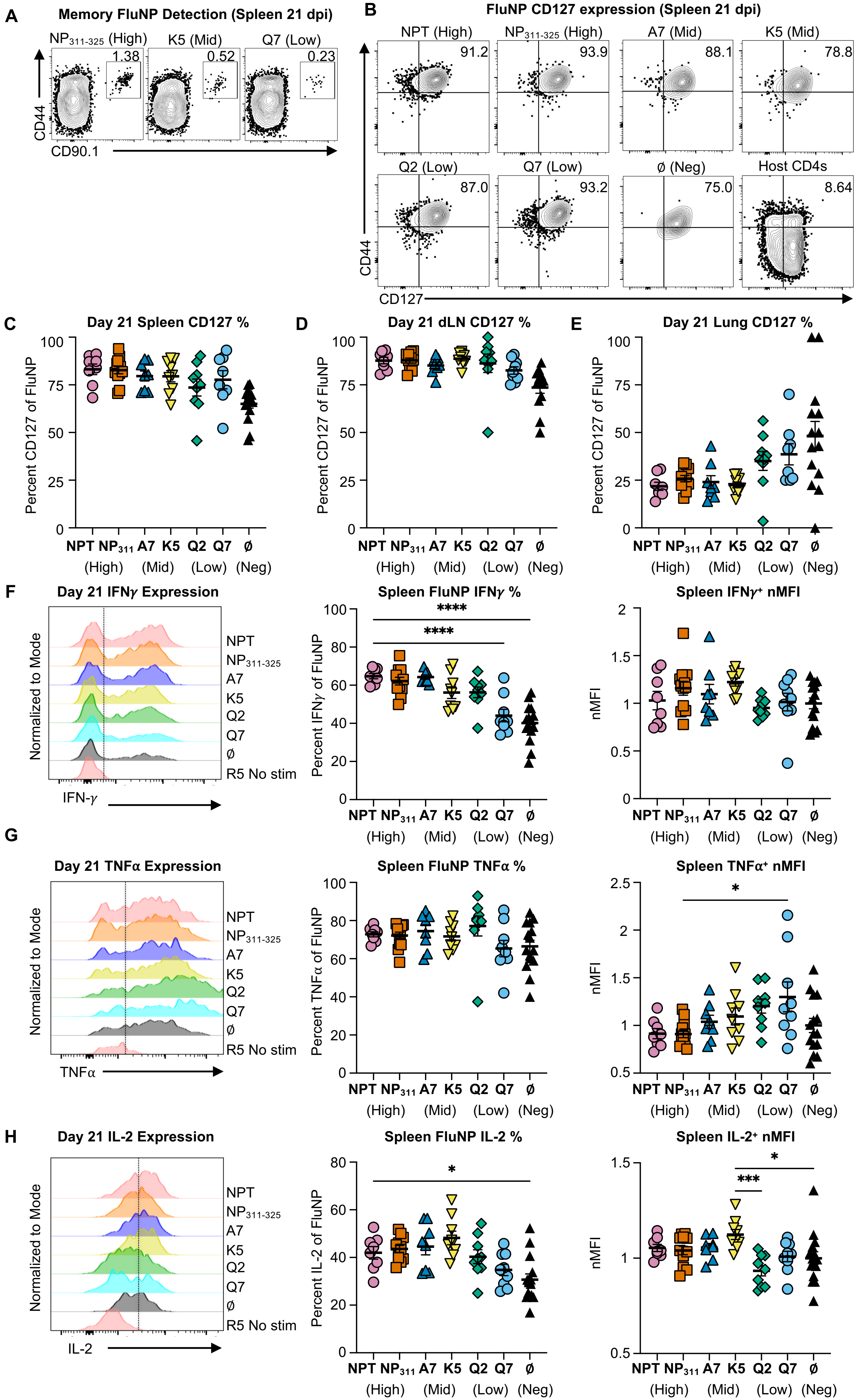


**Supplementary Figure 2. Related to Figure 2. Memory FluNP gating, phenotyping and cytokine production.**

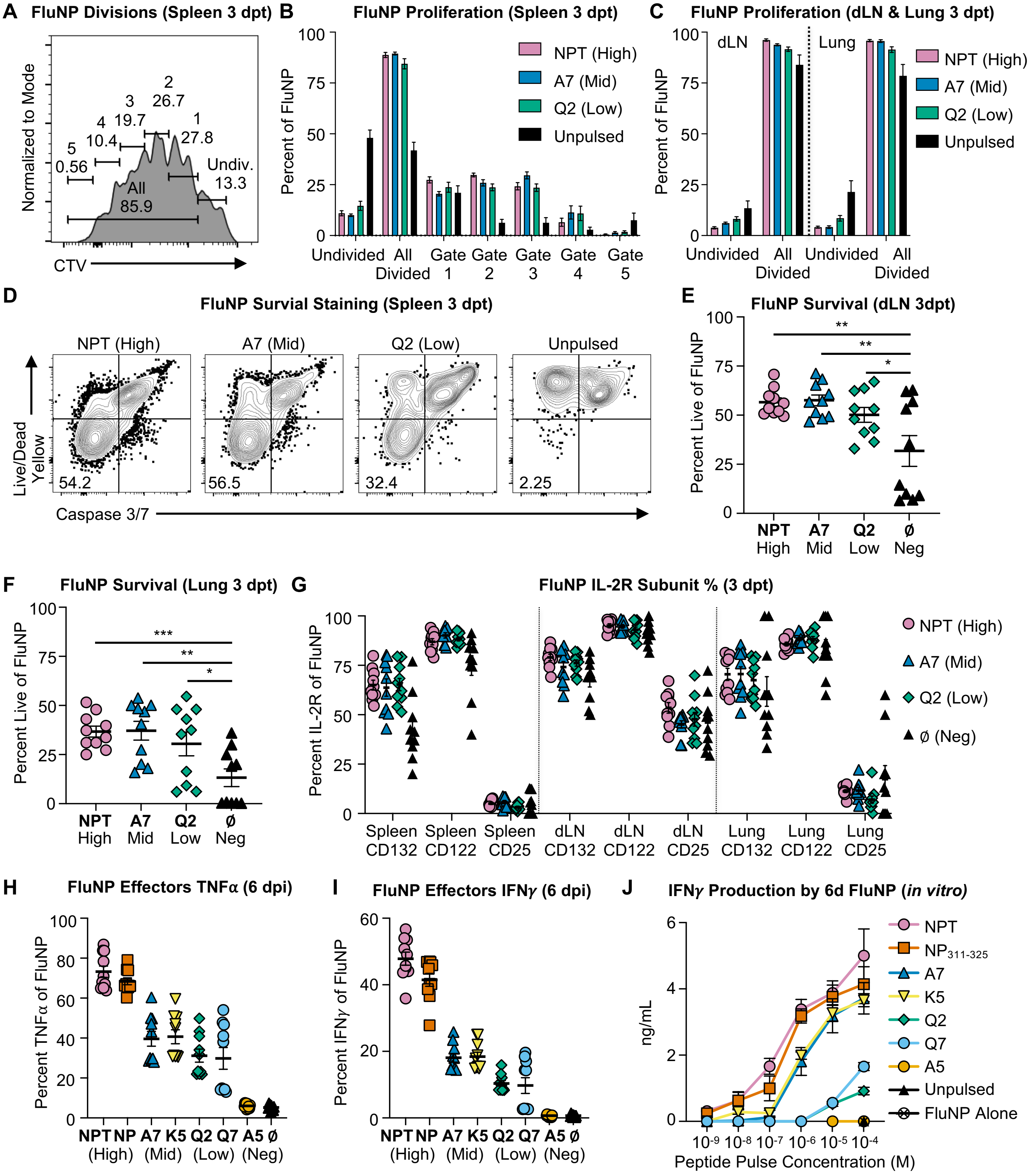


**Supplementary Figure 3. Related to Figure 3. Donor cell survival and proliferation and IL-2R subunit expression 3 days post transfer, day 6 cytokine production**

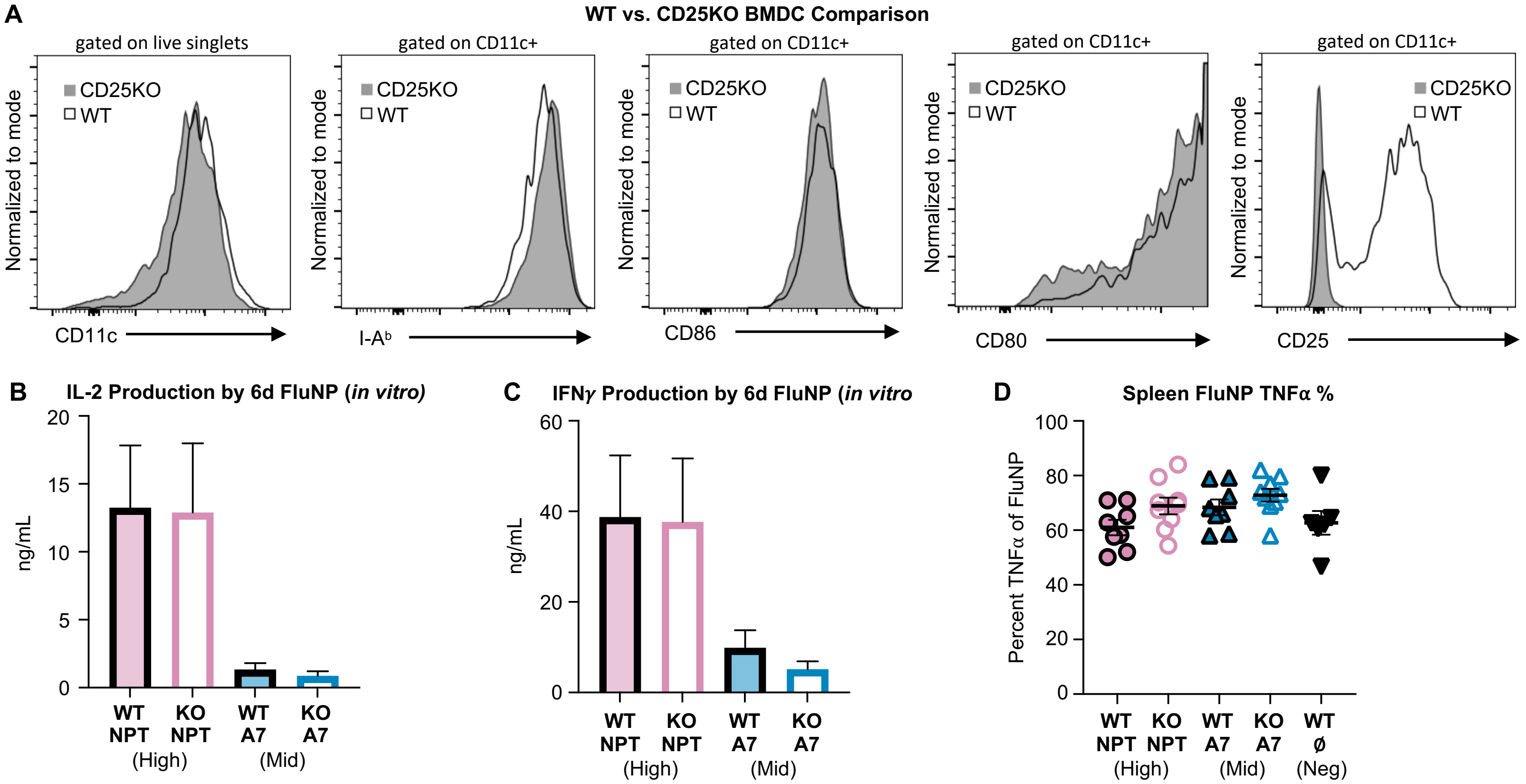


**Supplementary Figure 4. Characterization of WT and CD25KO APC, and their impact on FluNP effector cytokine production.**
